# Data-Driven Reduced Modeling of Recurrent Neural Networks

**DOI:** 10.1101/2025.10.13.681116

**Authors:** A. Marraffa, R. Krause, V. Mante, G. Haller

## Abstract

Artificial Recurrent Neural Networks (RNNs) are widely used in neuroscience to model the collective activity of neurons during behavioral tasks. The high dimensionality of their parameter and activity spaces, however, often make it challenging to infer and interpret the fundamental features of their dynamics.

In this study, we employ recent nonlinear dynamical system techniques to uncover the core dynamics of several RNNs used in contemporary neuroscience. Specifically, using a data-driven approach, we identify Spectral Submanifolds (SSMs), i.e., low-dimensional attracting invariant manifolds tangent to the eigenspaces of fixed points. The internal dynamics of SSMs serve as nonlinear models that reduce the dimensionality of the full RNNs by orders of magnitude.

Through low-dimensional, SSM-reduced models, we give mathematically precise definitions of line and ring attractors, which are intuitive concepts commonly used to explain decision-making and working memory. The new level of understanding of RNNs obtained from SSM reduction enables the interpretation of mathematically well-defined and robust structures in neuronal dynamics, leading to novel predictions about the neural computations underlying behavior.

## 1 Introduction

Behavior emerges from concerted activity across large populations of neurons. In the study of neural population activity, two primary questions emerge: How do neurons collectively create an internal representation of the external world based on the inputs they receive, and how do neural computations on these representations lead to behavior?

Recent advances in neuronal modeling and analysis have made it possible to address these questions, providing new insights into the nature of computations implemented at the level of neural populations. To establish a connection between the firing patterns of neural populations and low-dimensional, interpretable behavioral outputs, individual neurons can be interpreted as degrees of freedom within a dynamical system. Computations can then be understood in terms of the evolution of this dynamical system when driven by the inputs it receives from the senses or from other areas [1], [2]. Dynamical systems modeling naturally lends itself to analyses based on systems identification, whereby neural computations reflect the properties and interaction of low-dimensional, attracting structures, even when single-neuron responses are not directly interpretable.

Accumulating evidence from such approaches suggests that specific classes of dynamical motifs are consistently involved in cognitive and motor operations. For instance, saddle-point dynamics have been associated with decision-making processes [3], [4], and rotational dynamics have been proposed as a hallmark of population activity underlying movement preparation and execution [5] [6]. Some other heuristics are also helpful in characterizing observed dynamics in more complex tasks: “ring attractors” have been proposed as substrates for working memory [7], [8] [9] [10], [11], [12], and “line attractors” have been discussed for tasks that require the accumulation of evidence [13][14], [15], [16]. Importantly, more involved and structured behavioral tasks could be understood as the composition of such dynamical primitives [17], [18].

Identifying such dynamical motifs requires accurate models of neural dynamics. To this end, a wide range of methods has been developed in recent years to infer dynamics from population—level recordings. Many techniques rely on fitting a model of dynamics to the data (see [19] for an exception), but generally suffer from a tradeoff between complexity, expressive power, and the interpretability of the fitted models. At one extreme, machine—learning—based methods such as LFADS [20] and CEBRA [21] (see also [22], [23]) can be used to fit highly nonlinear, even chaotic dynamics, but often at the expense of mechanistic insight and predictive power. At the other end are models based on linear dynamical systems, either a single one [24], or switching [25], piecewise linear [26], or time-dependent linear systems [27]. Since the possible dynamics of linear systems are well understood, the resulting descriptions of dynamics are highly interpretable, but cannot capture intrinsically nonlinear behavior, such as coexisting attractors and chaotic dynamics.

Here, we propose a new approach to estimate neural dynamics based on a recent nonlinear model reduction technique that strikes a balance between the ability to capture intrinsically nonlinear dynamics and interpretability. Central to our approach is the recent theory of Spectral Submanifolds (SSMs)[28], which are ubiquitous and robust low-dimensional, attracting invariant manifolds that emanate from steady states with nondegenerate and nonresonant linearized spectra (see [29] for a general introduction, applications, and further references). The internal dynamics of SSMs serve as low-dimensional, explicit, polynomial models that explain the dominant slow nonlinear dynamics of high-dimensional dynamical systems. Based on these reduced models, dynamical motifs in neural networks can be analyzed and explained in a mathematically rigorous way.

We demonstrate the power of data—driven SSM—based modeling for neural dynamics using simulation output data from recurrent neural networks (RNNs). RNNs trained on specific behavioral tasks have emerged as a key tool to reproduce the features of neural dynamics in brain recordings and to generate hypotheses about the neural computations implemented by the underlying neural circuits [30], [31], [32]. However, obtaining a complete understanding of the dynamics of RNNs is often challenging, even when all their underlying parameters are fully known. Commonly used techniques for identifying dynamical motifs in RNNs rely on the search for fixed points and “slow points” [33], [13], [18], i.e., locations in neural state space where dynamics does not change or varies slowly. The geometry of slow points is largely conserved across different RNNs solving the same tasks and is therefore considered a useful heuristic of the computations they implement [34]. Further insights into computations can be obtained through linear approximations of the dynamics around the identified fixed points.

Such standard approaches to understanding RNN computations have several downsides, which are overcome by SSM-based modeling. First, slow points are typically found numerically based on subjectively chosen thresholds on the rate of change of neural dynamics. Second, linear approximations of non-linear systems are valid only only locally around fixed points, but not around slow points. Third, even when dynamics can be linearized, around hyperbolic fixed points the linear approximations are only valid in domains with simple, *linearizable* dynamics. These domains cannot contain multiple isolated fixed points, periodic orbits, quasiperiodic tori, chaotic attractors, or transitions among such invariant sets. In contrast, we show that data-driven algorithms to estimate low-dimensional SSM and their reduced dynamics ([35]) can provide a complete, yet simple description of both linear and nonlinear dynamics across all computationally relevant activity domains. The resulting mathematical descriptions of the global dynamics provide the basis for more precise hypotheses about the nature of neural computations in biological neural networks.

## 2 Results

We model recurrent neural networks with evolution rules of the form:

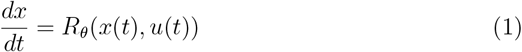

where *x*(*t*)∈ ℝ ^*N*^ is the network state variable, 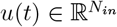 is the input vector and *R*_*θ*_ is a function depending on the parameter vector *θ* that defines the network dynamics.

Our approach to analyzing autonomous, high-dimensional RNN dynamics begins by assuming the existence of a known steady-state solution, which in practice can be identified as zeros of the right-hand-side of Eq. 1. Around such a steady state, we can deduce the existence of attracting invariant SSMs.

SSMs can be technically defined as the smoothest invariant manifolds that are tangent to dominant eigenspaces of steady states. The existence of these manifolds is ensured under nonresonance conditions on the spectrum of the linearized dynamical system at the steady state [29]. Once the existence of an SSM is established, it can be approximated through a polynomial expansion over the dominant spectral subspace used in its identification. While the existence of SSMs is inferred from the linearized spectrum at a steady state, these invariant manifolds do extend beyond the domain of linearization and hence can carry characteristically nonlinear dynamics. The internal dynamics of attracting SSMs serve as a mathematically exact reduced-order model with which nearby trajectories synchronize exponentially fast.

We employ the SSMLearn algorithm [35] to find explicit polynomial parametrizations for low-dimensional, attracting, invariant SSMs attached to steady-state solutions. SSMLearn then approximates the reduced polynomial ordinary differential equation (ODE) that governs the system evolution restricted to the SSM, providing a low-dimensional, polynomial model that captures the core dynamics of the original RNN.

A key advantage of this approach is the substantial reduction in model dimensionality, which converts a high-dimensional RNN into a low-dimensional polynomial ODE. Therefore, SSM-reduced models allow for the explicit identification of phase space structures, such as fixed points, limit cycles, and bifurcations, that determine the global asymptotic behavior of the network. To assess the effect of system parameters on dynamics 1, we can construct a slow manifold for each value of a given system parameter and view it as a section of a global slow manifold in the extended phase space that includes the system parameters as well. This, in turn, enables us to build parameter-dependent polynomial models for the SSM-reduced dynamics.

### 2.1 A Model for Context-Dependent Decision-Making

As a first example, we analyze the 100D RNN performing the context-dependent decision-making task introduced by [13], as described in the Methods section 5.1. This network receives as input two sensory cues, *s*_1_ and *s*_2_, whose values can be positive or negative, and two context cues, (*c*_1_, *c*_2_) with values in {(0, 1), (1, 0)}. The network’s output is expressed through a scalar readout function that will have the same sign of *s*_1_, if (*c*_1_, *c*_2_) = (1, 0), or *s*_2_, (*c*_1_, *c*_2_) = (0, 1).

The authors of [13] explain the network dynamics underlying flexible decisionmaking using the concept of a *line attractor*, which they view as a curve in phase space consisting of *slow points* [36], [13], [33]. Although not an attractor in a strict mathematical sense, the line attractor is a heuristic used to explain what is observed in the first principal component space: trajectories are attracted to this curve when sensory inputs are off and diverge from it once sensory inputs are turned on.

Within this 1D structure, [13] finds two stable fixed points that represent the two possible decisions in the task. The final state of the network, i.e., the fixed point to which it converges, is then argued to depend on the direction of perturbation away from the line attractor, determined by the direction and magnitude of the input vector (see Fig. 3 in [13] and Fig. 2 in [37]).

Here we revisit this interpretation through SSM theory, reinterpreting the RNN behavior as reduced dynamics on a 1D invariant slow manifold. We begin by analyzing the system under fixed context and sensory input, considering two conditions: the sensory input-off regime, corresponding to the cue period before sensory integration in animal experiments, and the sensory input-on regime, where active sensory processing occurs.

In both cases, we identify at least one fixed point in the system’s phase space, which ensures the existence of SSMs for the flow map. In the sensory input-on case, the network has an unstable fixed point and two stable fixed points, each corresponding to a distinct choice in the task. Importantly, these two stable fixed points are present for every input parameter value in the range explored in this experiment (see Fig. S2 in the Supplementary Materials). Linearization around the unstable fixed point reveals a single unstable eigenvalue, indicating a 1D unstable subspace. We use SSMLearn to find the expansion coefficients of the corresponding unstable manifold of the nonlinear system (Fig. 1, upper left and Fig. S1, right) up to order five and derive the associated reduced-order model

**Fig. 1:**
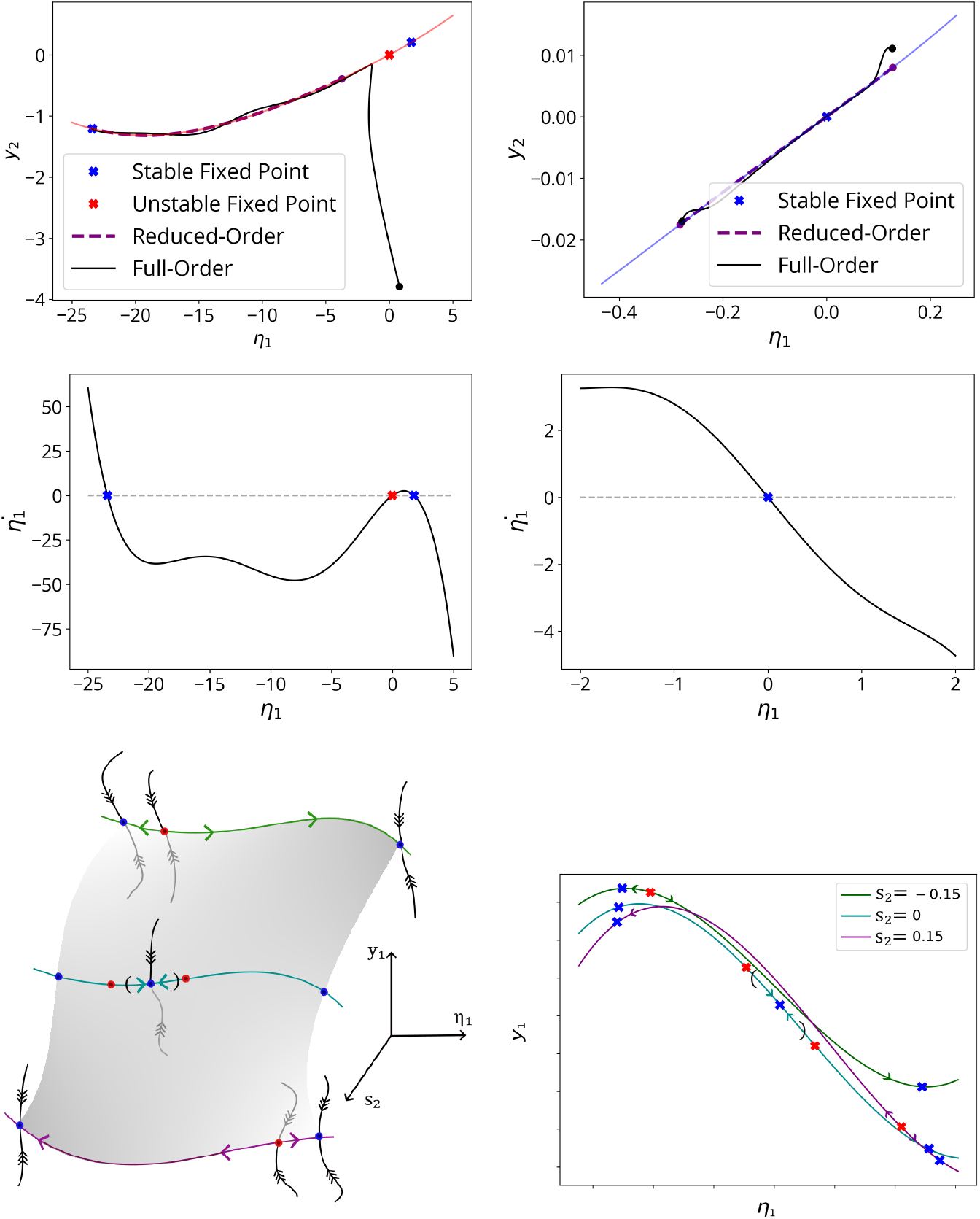
SSMs carrying the RNN reduced dynamics. (Upper left) Unstable manifold at order five in coordinates (*η*_1_, *y*_2_), where *η*_1_ parametrizes the spectral subspace *E*_1_ and *y*_*i*_ = *x*_*i*_ *−x*_*i*,0_ are the original RNN coordinates centered around the unstable fixed point. We depict a full-order test trajectory and the corresponding reduced trajectory on the SSM (after 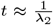, when transients have decayed, purple dot.) converging to the fixed point with the larger domain of attraction along the unstable manifold. Black dots correspond to initial conditions. The fixed points in the plot (crosses) are the fixed points of the full model, lying on the unstable manifold and correctly reproduced by the reduced-order model. The Manifold Fitting Error (MFE), the mean distance between observed trajectories and trajectories projected on the manifold, has magnitude *≈* 0.007, and the Normal Mean Trajectory Error (NMTE), the mean reconstruction error of the reduced model, is *≈* 0.03. (Upper right) The slowest SSM at order three and test full-order and reduced-order trajectories plotted in coordinates (*η*_1_, *y*_2_), with *y*_*i*_ now centered around the stable fixed point. The MTE and NMTE have magnitude *≈* 0.05. (Center) Graphs of the right-hand sides of the reduced-order models: on the 1D unstable manifold (left) and on the 1D slowest SSM (right), both at order five. On the left plot, the unstable fixed point (red cross) serves as a boundary for the domains of attraction of the two stable fixed points (blue crosses) on the unstable manifold. On the right plot, we observe convergence to the stable fixed point. (Bottom) Sketch of the robust (normally hyperbolic) 2D invariant manifold (*slow manifold)* in the extended phase space constructed including the *s*_2_-parameter direction (left), in coordinates (*η*_1_, *s*_2_, *y*_1_), where *η*_1_ and *y*_1_ are as above, and in coordinates (*η*_1_, *y*_1_) for selected *s*_2_ values (right). We see how the 1D latent dynamics underlying flexible decision-making change when varying the relevant sensory input *s*_2_ from negative to positive values, passing through 0, resulting in a change in the preferred choice for the network. When *s*_2_ = 0, there is a stable fixed point that attracts initial conditions around zero and, locally, the slow manifold coincides with its 1D slowest SSM (in round brackets).

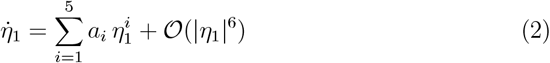

whose right-hand side is plotted in Fig. 1 (center left). The SSM-reduced model (2) at this order of approximation enables accurate prediction of the full RNN dynamics from simulations of the reduced model.

In fact, the 1D model (2) is sufficient to reproduce the RNN behavior during decision making. On the unstable manifold, the size of the domains of attraction of the two stable fixed points determines the global phase space geometry: random initial conditions are more likely to evolve toward the fixed point associated with the larger domain of attraction, which corresponds to the correct choice in the task. We estimate these domains of attraction in the full phase space using Finite-Time Lyapunov Exponent (FTLE) analysis in the Supplementary Materials S3. FTLE ridges identify regions of phase space where initial conditions close to each other have a high rate of separation. In this context, these initial conditions mark the boundaries of the domains of attraction of the two fixed points (see Fig. S4).

In the sensory input-off case, the dynamics in a small neighborhood of the origin (where initial conditions are set in the experiments) are governed by a stable fixed point with one slowest eigenmode in its linearized dynamics. Using SSMLearn, we find the expansion coefficients the corresponding 1D slowest SSM (see Fig. 1, upper right). Full details and the corresponding equations for the reduced dynamics, whose graph is shown in Fig. 1 (center right), are provided in the Supplementary Materials S1.

These results lead to a re-interpretation of the line attractor in [13]. The set of slow points identified in [13] during the sensory input off-period corresponds to the slowest SSM anchored to the stable fixed point. When sensory inputs are off, this manifold attracts nearby trajectories and carries stable, near-equilibrium dynamics. These dynamics are perceived as slow once the trajectories are close enough to the stable fixed point, to which they converge in infinite time.

When sensory inputs are activated, the dynamics transition: trajectories converge to the decision-related stable fixed point along the 1D unstable manifold of a now unstable fixed point (in Fig. 3 of [13], trajectories labeled as sensory input on). The reduced dynamics restricted to this SSM exhibit a bistable structure, wherein the unstable fixed point acts as a boundary separating the domains of attraction of the two stable fixed points.

Importantly, as depicted in Fig. 1 (bottom), a 1D slow manifold is present throughout the explored range of sensory inputs and carries the relevant dynamics throughout the decision-making process. Specifically, when sensory inputs are zero, the slow manifold coincides locally with the slowest SSM attached to a stable fixed point. As the relevant sensory input *s*_*i*_ (with *i* = 1 for context 1 and *i* = 2 for context 2) is varied from zero, we retrieve the slow manifold as the unstable manifold of a (different for different *s*_*i*_ signs) unstable fixed point that separates the basins of attraction of the two stable equilibria (see Fig. S2 and discussion therein in the Supplementary Materials).

As we show in the Supplementary Methods S4, we can assess the robustness of the 1D slow manifold as sensory inputs change, and we know that its geometry and associated reduced dynamics deform smoothly. In other words, in the extended phase space (constructed including the parameter axis, as in Fig. 1, bottom left), we find a 2D, at least *C*^1^ attracting slow manifold that carries the latent dynamics underlying flexible decision-making. Around persisting fixed points, we can explicitly determine the dependencies on parameters through parameter-dependent polynomial expansions on their SSMs (see Supplementary Materials S2 for a complete discussion).

Our analysis also clarifies the role of the input vector and the recurrent dynamics in determining the outcome of the decision-making task, allowing us in particular to distinguish among the three hypotheses proposed by [37]. Notably, we find that the direction of the input vector in the reduced dynamics does not, by itself, determine the result of sensory integration (see Fig. 2). Moreover, as illustrated in Figs. 1 (bottom), 2, and S3, the geometry of the SMMs remains largely constant across context and sensory inputs. As the input vector changes orientation, the tangent spaces of the SSMs do not align with it to facilitate integration. Instead, convergence is determined by the location of initial conditions relative to the two domains of attraction. The width of these domains (and the location of the separating unstable fixed point on the slow manifold) varies parametrically with the relevant sensory input (or with its mean in the noisy case) to achieve the desired outcome. In other words, mean input vectors are parameters that determine the geometry of the phase space of the RNN.

**Fig. 2:**
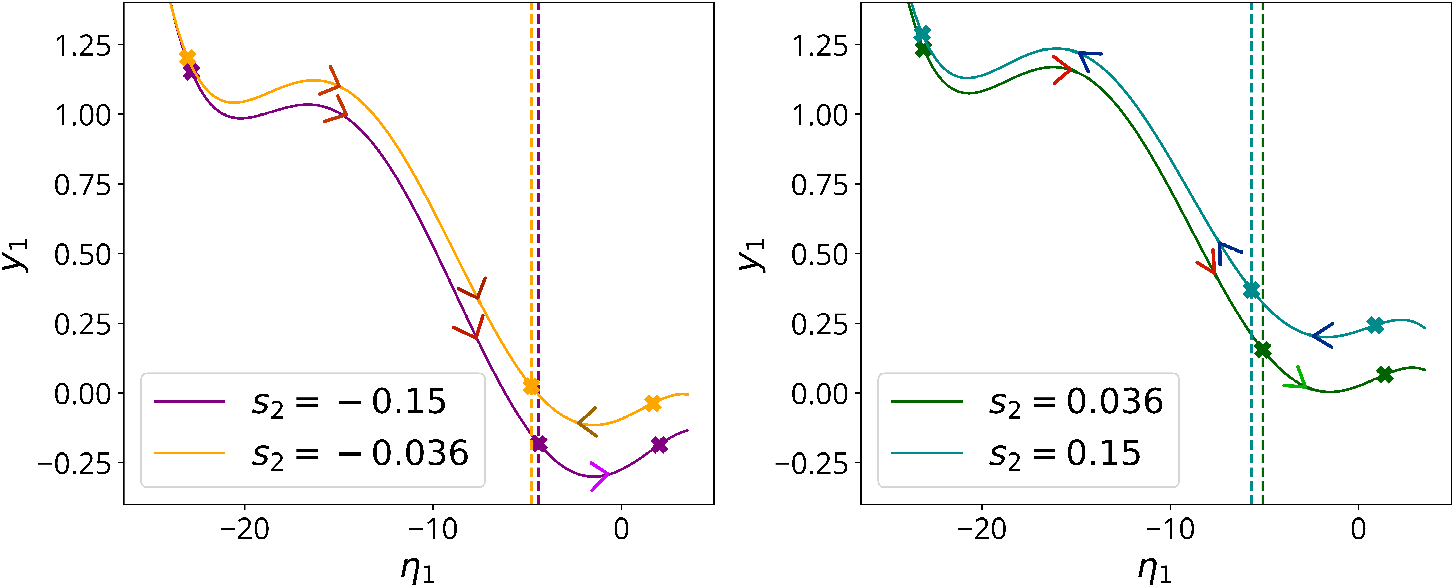
One-dimensional slow manifolds (unstable manifolds) of the system for different values of the sensory input parameter *s*_2_ (selected by context 2). The fixed point (coloured crosses) the network has learned to converge to depends on the sign of *s*_2_. The way the RNN realizes this input-output relationship is changing location to the unstable fixed point (central cross) along the slow manifold depending on *s*_2_: the unstable fixed point separates the domains of attraction of the two stable fixed points in the reduced dynamics, so its location (dashed line) determines the asymptotic behavior of trajectories, given their initial conditions. The crosses correspond to the fixed points, the two stable one divided by the unstable one. We display the arrows indicating effect of the input vector on the reduced dynamics (projection onto the SSM). This picture shows that the projection of the input vector (or its mean, in the noisy case) on the reduced dynamics does not determine the asymptotic behavior (choice) of the RNN, as we can see that, for example, it would lead to the wrong direction for *s*_2_ = 0.036 (red arrows). However, whenever the perturbed initial conditions lie within the correct domain of attraction, the convergence to the correct choice is guaranteed (see also S3).

In summary, based on the above findings we propose an alternative to the *line attractor interpretation* of these RNN, namely a mathematically rigorous 1D *slow manifold explanation*, and provide a method to systematically find these slow manifolds emanating from known steady-state solutions.

More generally, 1D slow manifolds may play a central role in decision-making tasks, not only in RNNs but also in biological neural systems. For instance, [38] shows that a linear Vanilla RNN trained on mouse behavioral data from a sensory discrimination task learns a 1D slowest mode during training. This observation suggests that SSM theory has the potential to aid the analysis of neural computations in both artificial and biological settings. Importantly, the derivation of SSMs extends to cases where activity is corrupted by noise. We discuss the effects of adding noise to the RNN dynamics in the Supplementary Materials S1, where we treat uniformly bounded Gaussian noise as a small-amplitude, time-dependent perturbation to the RNN studied so far.

### 2.2 A Model for Oscillations with Input-Dependent Frequency

Our second example, the 100D *Sine Wave Generator* RNN in [39] (see Method section 5.2), is trained to produce a sinusoidal scalar readout *z*(*t*), with oscillation frequency selected by a scalar input *u* (see Eq. (14)).

In the phase space of the constant-input RNN, there is an unstable fixed point with two complex-conjugate unstable eigenvalues widely separated in their real parts from the stable ones. As a consequence, a 2D slow SSM exists in this problem and coincides with the unstable manifold of the fixed point. This SSM carries the observable oscillatory dynamics of the system. We find the SSM equations and its reduced dynamics with SSMLearn up to order three and six, which are the lowest expansion orders that give low invariance error (distance of trajectories from the SSM) and prediction error (distance between full and reduced trajectories), respectively. Specifically, the SSM-reduced dynamics takes the form

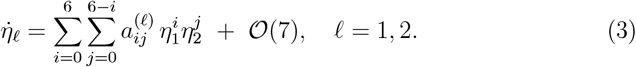

Analyzing the SSM-reduced dynamics (3), we find a globally attracting limit cycle that is responsible for the periodic oscillations with the desired frequency. This type of dynamics is exactly the simplest possible realization of a dynamical system that oscillates with a single frequency.

The unstable manifold, the limit cycle, and full and reduced trajectories are plotted in Fig. 3 (upper left), while contour lines of the vector field defining the reduced-order model found through SSMLearn at order seven, projected to the unstable subspace *E*_2_ can be seen in Fig. 3 (bottom left). Hence, we obtain a 2D invariant manifold and reduced-order model which provide a complete and detailed description of the dynamics of this high-dimensional, oscillatory RNN. When we choose an input *u* with frequency *f* = 1.9 Hz, the frequency of the resulting limit cycle captured by the SSM-reduced model is *f*_*RED*_ = 1.92 Hz, which is very close to the measured frequency *f*_*F ULL*_ = 1.91 Hz in the full RNN model.

**Fig. 3:**
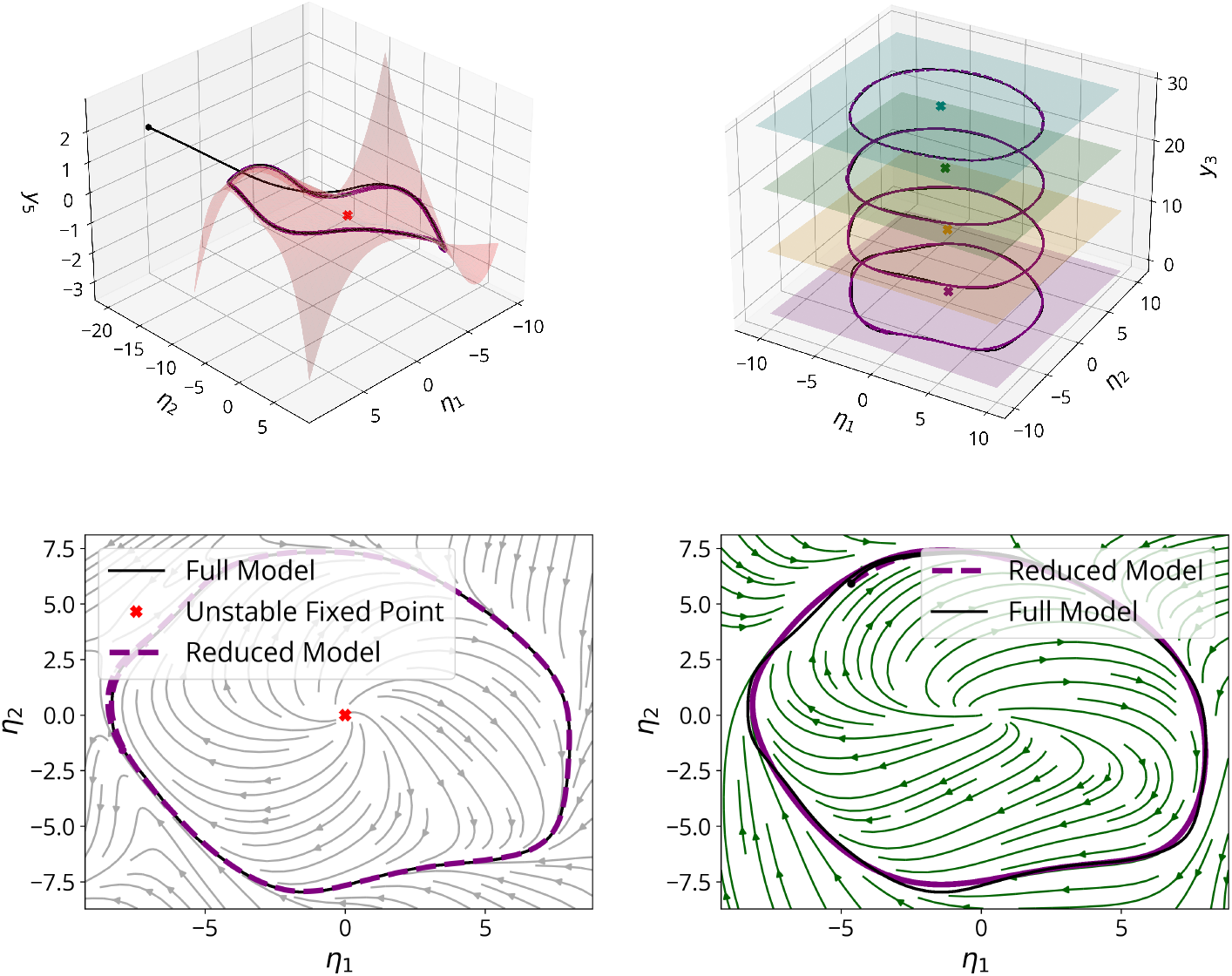
SSM carrying the core RNN dynamics and parameter-dependent SSMs with their reduced dynamics projected to the spectral subspaces. (Upper left) Unstable manifold at order three in coordinates (*η*_1_, *η*_2_, *y*_5_), where (*η*_1_, *η*_2_) are the coordinates of the unstable subspace. The stable limit cycle on the unstable manifold is globally attracting: in the picture, one full and one reduced trajectory are converging to the stable limit cycle. The MFE and the NMTE calculated on test trajectories have magnitude *≈* 0.02 and *≈* 0.5, respectively. (Bottom left) Projections onto the unstable subspace of the streamlines of the right-hand side of the reduced-order model on the two-dimensional unstable manifold up to order six, with projected full and reduced trajectories. (Upper right) Two-dimensional unstable manifolds for different values of the parameter *u* (and corresponding output frequency values *f*) plotted in the three-dimensional space with coordinates (*η*_1_, *η*_2_, *y*_5_). The manifolds are shifted on the *y*_5_-axis for visualization purposes. (Bottom right) Projection onto the unstable subspace of the streamlines of the right-hand side of the reduced-order model on the two-dimensional unstable manifold up to order six for parameter *u* corresponding to a frequency of 3.9 Hz, with projected full and reduced trajectories.

To analyze the dependence of the limit cycle frequency on the input parameter *u*, we recall that unstable manifolds depend smoothly on parameters, as long as their underlying unstable spectral subspaces persist under the variation of those parameters (see, for example, [40]). As a consequence, we have a smooth, *u*-dependent family of 2D unstable manifolds in the phase space, as illustrated in Fig. 3 (upper right).

Along members of this SSM family, we have an explicit, *u*-dependent polynomial expansion for a 2D, parametric, SSM-reduced model of the form

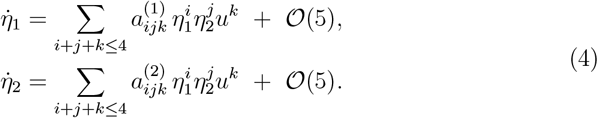

The limit cycle frequencies in the parametrized model (4) agree with the measured frequencies of the full trajectories up to an error of 10^*−*1^.

This example illustrates how our reduction approach can uncover the specific solution that the training process of an RNN selects from many potential solutions to perform a given task. Rather than viewing RNNs as black box systems that replicate some behavior, SSM-reduced modeling uncovers the low-dimensional, interpretable core dynamics 3 of RNNs. This in turn enables us to identify a very specific dynamical structure, a limit cycle, that is directly responsible for producing the output that the RNN was trained to produce.

### 2.3 A Model for a Memory-Pro Task

As our third example, we now describe the SSM-based reduction of the multitasking RNN in [18] performing a Memory-Pro Task described in the Method section 5.3. This RNN receives as input a fixation input *u*_1_ with values in {0, 1}, and two 2D angular sensory inputs *u*_2_ = cos *θ*_1_, *u*_3_ = sin *θ*_1_, *u*_4_ = cos *θ*_2_, and *u*_5_ = sin *θ*_2_. In the setting considered here, the network has to respond according to the angle indicated by the first stimulus when the fixation input is off (*u*_1_ = 0). This is expressed through a three-dimensional readout vector **z**: *z*_1_ = *u*_1_ (fixation) and *z*_2_ = cos *θ*_1_,*z*_3_ = sin *θ*_1_.

The authors of [18] interpret the network dynamics underlying the memory period within the task through a 1D structure composed of slow points [36, 13, 33] located along a circle, which they call a *ring attractor*. In this structure, the position that a trajectory reaches during the memory period is interpreted as the persistent pattern that the network maintains over time, encoding the correct stimulus it has to respond to when the fixation input is off.

For this reason, we focus on the input parameter setting corresponding to the memory and context periods, when fixation and context inputs are on and sensory inputs are off. For this configuration, we seek to clarify what dynamical structure is behind what has been empirically labeled as a ring attractor. Such clarification is necessary, as the attractor topology closest to a ring would be that of a torus. There is, however, no observational evidence for sustained quasiperiodic behavior, which would be the hallmark of a toroidal attractor.

Our analysis reveals an unstable fixed point with two unstable, real eigenvalues and an asymptotically stable fixed point. Based on the spectrum of the linearized dynamics around the unstable fixed point, we conclude that the 2D SSM embodied by the unstable manifold of the unstable fixed point will carry the observable core dynamics once the transients have died out.

We find the unstable manifold using SSMLearn up to polynomial order three (see Fig. 4, upper left). The reduced-order model on this SSM is of the form

**Fig. 4:**
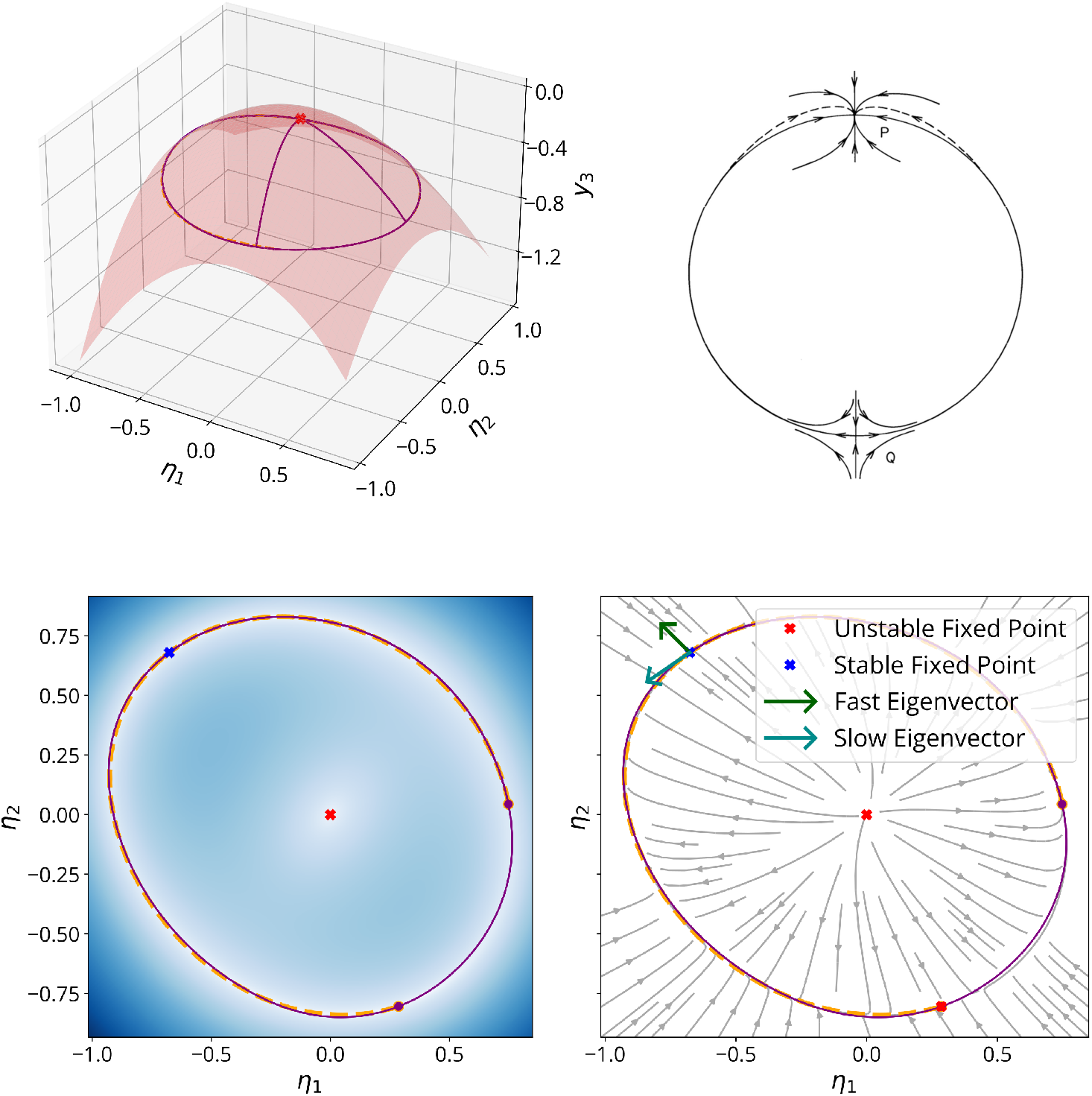
The heteroclinic orbit on the 2D SSM. (Upper left) The two-dimensional unstable manifold attached to the unstable fixed point carries the reduced dynamics. Trajectories (full model trajectories in purple and reduced model in orange) disposed on a circular curve converge to the stable fixed point in infinite time. (Bottom left) Norm of the right-hand side of the reduced model on the unstable manifold plotted on the unstable subspace with coordinates (*η*_1_, *η*_2_), with darker blues corresponding to larger norm values. The MFE and the NMTE calculated on test trajectories have magnitude 10^*−*2^ and 10^*−*1^, respectively. (Bottom right) Heteroclinic orbits connecting the unstable fixed point on the bottom right and the stable fixed point on the upper left, projected onto the unstable subspace with coordinates (*η*_1_, *η*_2_). Streamlines of the right-hand side of the reduced model are projected onto the unstable subspace (in gray). The structure is normally hyperbolic, as we can see from the directions of the slow and the fast eigenvectors of the linearization around the stable fixed point of the full model. (Upper right) Pictorial representation of the normally hyperbolic heteroclinic structure, as an example of normally hyperbolic invariant manifold in [41]. The unstable manifold of the unstable fixed point Q coincides with the slow stable manifold of the stable fixed point P, and the rate of normal attraction to P is bigger than the rate of tangential compression.

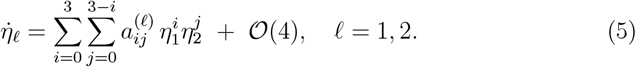

Solving for the zeros of the right-hand side of (5), we find an additional unstable fixed point. This fixed point is connected to the stable fixed point on the SSM via a pair of heteroclinic orbits that form a structurally stable heteroclinic loop (see Fig. 4, bottom). The structural stability (i.e., smooth persistence under perturbations of the RNN) of this heteroclinic loop follows from the fact that it is normally hyperbolic, i.e., the attraction rates normal to it dominate the attraction rates inside it ([41], [42]).

This strict result from the SSM-reduced model (5) clarifies that the true phase space structure behind the empirically observed slow circular structure (ring attractor) is an attracting, 1D invariant manifold. Diffeomorphic to a circle, this manifold is formed by a structurally stable heteroclinic loop between two fixed points inside a 2D slow SSM.

The structural stability of normally hyperbolic heteroclinic loops explains the ubiquitous presence of slow, circular attractors in RNNs performing working memory tasks (see [7], [8] [9] [10]). If, in fact, working memory corresponds to the maintenance of persistent neuronal firing patterns, represented in our framework as fixed points in phase space, it is tempting to conclude (see [16, 11, 12, 14, 15, 43, 44, 45]) that encoding a continuous variable would require a continuum of such fixed points, each parameterized by a possible value of the variable.

A continuum of fixed points, however, is inherently non-robust and can be destroyed by small perturbations to the neuronal dynamics. Instead, robust circular attractors that are not limit cycles are necessarily formed by chains of heteroclinic connections among fixed points. Here, we have observed the simplest such chain, one that is formed just by two heteroclinic orbits.

## 3 Discussion

We have shown how analyzing RNNs based on recent results from nonlinear dynamical systems theory reveals the core dynamics of those networks via considerably lower-dimensional, polynomial models on attracting SSMs. Our results firm up previous, mostly heuristic interpretations of RNN dynamics. Specifically, the low dimension of the resulting SSM-reduced models allows us to develop a detailed mathematical understanding of the global RNN behavior. We describe these dynamics in terms of attracting invariant manifolds, the invariant sets they carry, and the transitions among those sets.

In particular, we have obtained a one-dimensional (1D) parameter-dependent model on a *slow manifold* for the 100D, context-dependent decision-making RNN in [13]. In a similar, data—driven fashion, we have constructed 2D models on slow SSMs for the 100D sine wave generator RNN in [39], and the 256D multitasking RNN in [18] performing a memory-pro task. As opposed to slow manifolds arising in geometric singular perturbation studies (see [46]), SSMs exist without the assumption of a global time scale separation that divides the phase space into slow and fast variables. Therefore, SSMs exist in a more general setting in which the core dynamics settle on a low-dimensional manifold once the faster transient dynamics have died out.

Specifically, for the RNN performing the context-dependent discrimination task, we have explained the “line attractor” observed by [13] as a robust, parameter-dependent 1D slow manifold. For every configuration of the input parameters, we have identified the corresponding 1D slow manifold (as a section of the parameter-dependent slow manifold in the extended phase space) that captures the reduced dynamics of the full RNN. The resulting behavior following sensory integration arises from the dynamics on a 1D unstable manifold, along which the system converges to the stable fixed point associated with the correct choice.

We have also explained the flexibility of the decision-making process in terms of parameter-dependent SSMs and reduced models. In particular, context and sensory inputs determine which of the two fixed points on the 1D unstable manifold has the larger domain of attraction.

For the sine-wave generator RNN in [39], we have uncovered the dynamics responsible for the input-dependent sinusoidal readout with a simple 2D model on a 2D unstable manifold carrying an attracting periodic orbit. This explains how the frequency of this orbit varies with the scalar input through a parameter-dependent model on the parameter-dependent unstable manifold.

Finally, we have found that the empirically described “ring attractor” in the memory-pro task in [18] is a normally hyperbolic invariant slow manifold that coincides with two heteroclinic orbits connecting a stable and an unstable fixed point on a 2D SSM. The stable fixed point on the heteroclinic circle attracts nearby trajectories, and the heteroclinic structure carries slow, converging dynamics.

The relationship between RNNs trained to perform specific readout functions and the actual computations carried out by neurons is unclear, as many solutions are available for a high-dimensional system to produce a low-dimensional readout. However, an accurate description of the RNNs’ phase space geometry enables the generation and verification of hypotheses on the dynamical systems underlying the observed dynamics in the data (see [30], [47]). Interestingly, in the case of the 100D network trained for a binary choice task (context-dependent decisionmaking), the solution is a simple 1D bistable system. Similarly, we have found that a limit cycle with input-dependent frequency is responsible for the input-tuned oscillations of the sine-wave generator network. Moreover, the heteroclinic structure explaining the memory-pro task is a robust manifold that appears to be the common mechanism shared by all RNNs trained to reproduce short-term memory mechanisms [7], [8].

The data—driven SSM—reduction approach used here is general enough to be applied to RNNs performing more complex tasks, whether or not they are related to neural computations. Indeed, RNNs are high-dimensional, nonlinear dynamical systems, whose use extends beyond neuroscience to describe any system with rich dynamics. We rely on simulations of RNN trajectories to find the reduced models of their dynamics through SSMLearn [35]. This is a fully data-driven algorithm that is applicable to systems whose equations of motion are either unknown or do not display any particular symmetry. Our only assumption is that the RNN has at least one fixed point whose location is at least approximately known. Importantly, the dimension of the SSM—reduced model (typically one or two) is independent of the RNN dimension for generic parameter choices. Therefore, even for more complex RNNs not considered in this study, the potential for dimensionality reduction using SSMs is virtually unlimited.

The SSM theory was originally developed for model reduction in mechanical engineering, to reduce the dimensionality of high-dimensional mechanical systems, first in an equation-driven fashion [48], and later extended to fully data-driven problems [35]. However, neural systems are inherently different from mechanical systems, which generally have well-understood governing equations that are consistent with observations. For this reason, the application of SSM theory to neural data is promising but not straightforward. Recent extensions of SSM theory cover forced systems [49] and random systems with small, uniformly bounded noise [50]. Therefore, a starting point for SSM reduction carried out directly on experimental neural data will require a feasible model for noise in neuronal dynamics.

## Methods

### 4 The Theory of Spectral Submanifolds

To identify robust, smooth, and unique invariant manifolds in high-dimensional Vanilla RNNs, we employ the theory of SSMs, first formulated by Haller and Ponsioen in [28] to find invariant manifolds in high-dimensional nonlinear systems around an attracting steady-state solution. The reader can consult [29] for a complete and detailed introduction to the theory and its applications.

SSM theory is based on a fundamental observation about linear dynamical systems of the form

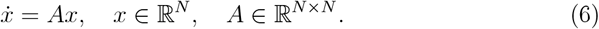

In this setting, the eigenspaces of *A*, as well as their algebraic direct sums, are invariant under the flow map and therefore constitute linear and smooth invariant manifolds of the system. We refer to any subspace of the form *E* = *E*_1_ ⊕ … ⊕ *E*_*k*_, where each *E*_*j*_ is the eigenspace associated with a particular eigenvalue *λ*_*j*_, as a *spectral subspace*. Important examples include the stable, center, and unstable subspaces, spanned respectively by the generalized eigenvectors corresponding to eigenvalues with negative, zero, and positive real parts.

From the eigendecomposition of *A*, we can always find solutions to the linear equations (6) and gather them together to construct infinitely many invariant manifolds tangent to each spectral subspace at the fixed point at the origin. Among these, the spectral subspaces are the smoothest (analytic), whereas all others have limited smoothness governed by the ratios of the real parts of the eigenvalues of *A*.

SSM theory exploits the structure of linear systems and uses linearization theorems to infer the existence, smoothness, and uniqueness of invariant manifolds, SSMs, in nonlinear systems around a fixed point. Although the existence of these manifolds is obtained from linearization, they extend over the domain of linearizability of the system and can carry nonlinear reduced dynamics. We look for SSMs as smooth invariant graphs tangent to the spectral subspaces of the linearized dynamics which, in the general case, are not invariant manifolds of the system themselves and hence not suited for model reduction (see [51] for further discussion).

In particular, consider the nonlinear autonomous system:

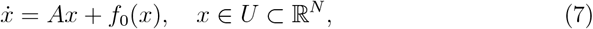

where *f*_0_ ∈ *C*^*r*^ for some *r* ∈ ℕ ∪ {∞, *a*} (where *C*^*a*^ indicates that the function is analytic) and satisfies *f*_0_(*x*) = *O* (∥*x*∥^2^) as *x* → 0. The fixed point at the origin is assumed to be hyperbolic and attracting, meaning that all eigenvalues of *A* have strictly negative real parts.

Given a spectral subspace *E* for the linearized dynamics, we define its *spectral quotient* as the integer ratio

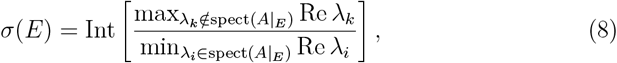

which compares the fastest decay rate outside *E* to the slowest decay rate within *E*. We can then define a *external nonresonance condition* for E (see [28]): no low-order (2 ≤ *m* ≤ *sigma*(*E*)) integer linear combination of eigenvalues within *E* should coincide with an eigenvalue outside of *E*.

Under the above assumptions, for a nonresonant spectral subspace *E* of the linearized dynamics of system (7) there exists an invariant manifold, *W*_*E*_(0), tangent to *E* at the origin. This manifold, the SSM associated with *E*, is the smoothest among all invariant manifolds tangent to *E*, and unique in the smoothest class *C*^*σ*(*E*)+1^. Furthermore, *W*_*E*_(0) inherits smooth dependence on coordinates and parameters from the right-hand side of (7).

In cases where the fixed point is unstable but has a nonempty stable manifold, a similar result holds, provided stronger nonresonance conditions are satisfied, specifically, that no eigenvalue of *A* is a nonnegative integer linear combination (with *m* ≥ 2) of all eigenvalues. Under these assumptions, one still obtains an invariant SSM, *W*_*E*_(0), that is unique in its smoothness class and tangent to the spectral subspace, as shown in [52].

In summary, attached to a hyperbolic fixed point with a nonresonant spectral subspace of the linearized dynamics, there is a unique invariant manifold, the SSM, distinguished by its highest smoothness. Since the SSM *W*_*E*_(0) is tangent to its underlying spectral subspace *E* at the fixed point, it is convenient to locally represent it as a graph over that spectral subspace.

As an example, let 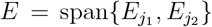 be a 2D spectral subspace for *A* in (7), with *f*_0_ ∈ *C*^*a*^ (analytic), for which the external nonresonance conditions are met. We assume that we have chosen a basis such that *E* is parameterized by the reduced coordinates (*η*_1_, *η*_2_), and the full coordinate system is given by *x* = (*η*_1_, *η*_2_, *y*_1_, …, *y*_*N* 2_). We look for the Taylor expansion of the SSM, *W*_*E*_(0), expressed as a graph over *E*

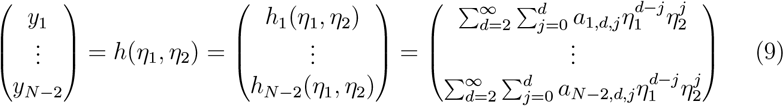

Since *W*_*E*_(0) is invariant, we can write the *invariance PDE*

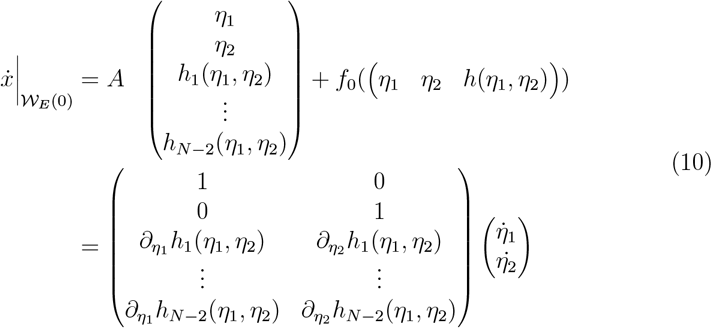

with boundary conditions at zero *h*(0) = *Dh*(0) = 0. Substitution of the Taylor expansion (9) into (10) yields a family of linear equations for the Taylor coefficients *a*_*i,j*_, which can be solved recursively under the asumed nonresonance conditions.

If the right-hand-side of (7) is not analytic but *C*^*r*^, *r* ∈ ℕ ∪{∞}, increasing the order of the expansion will not necessarily improve the accuracy of the approximation, even on smaller domains. If *f*_0_ is *C*^*r*^ with *r* finite, its expansion would not exist to orders higher than *r*. One should choose accurately the order to truncate the expansion, taking care not to exceed the order at which the approximation ceases to improve on the desired domain.

This theoretical basis for explicitly constructing invariant manifolds over arbitrary non-resonant spectral subspaces of the linearized system can be conveniently used for the reduction of high-dimensional nonlinear systems like RNNs. In particular, as a model for the slow, dominant dynamics, we can choose a *slow spectral subspace E* such that *E* = {*E*_1_ ⊕*E*_2_⊕ … ⊕ *E*_*q*_} is the direct sum of the *q* slowest eigenspaces whose corresponding eigenvalues have the largest real parts (they represent the lowest decaying dynamics in the linearized system if the fixed point is stable but also growing dynamics if the fixed point is unstable).

We point out that the dimensional parameter *q* in the choice of the slow spectral subspace *E* is a free parameter. It can be chosen so that the slow SSM becomes the low-dimensional attractor that prevails in the available data. For this purpose, we can examine the largest spectral gaps (difference in the real parts) between consecutive (ordered) eigenvalues and pick a slow spectral subspace that is separated from the others on a time scale suitable for the system under study, or simply the slow subspace corresponding to the largest gap.

Back to our example, once a functional expression for the SSM is obtained, we can find the reduced dynamics on the SSM by substituting the *y* variables with 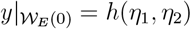

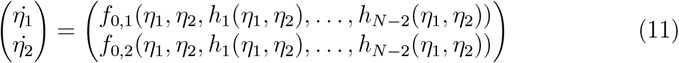

thus obtaining an explicit polynomial expansion for the 2D reduced-order model on the manifold. Since we have enslaved all the other dynamical variables to *η*_1_ and *η*_2_ through the SSM equation (9), we can retrieve the dynamics of the full system by inserting the solutions of the reduced equations (11) into the SSM equation.

The latest version of the algorithm that performs these equation-driven calculations is available in the open source MATLAB live script package SSMTool https://github.com/haller-group/SSMTool-2.4 (see [53]).

#### 4.1 Parameter-dependent and time-dependent models on SSMs

To understand the input-output relationship in Vanilla RNNs that resemble the sensory input-behavioral output connections in the animal tasks, we must consider how parameter changes affect the dynamics of the RNNs and their SSMs.

In the absence of noise or other types of explicit time dependence (which we may add later as perturbations), the input vector collects the parameters on which the system depends (see the discussion in section 5.1). Although altering sensory inputs and context inputs (see [13, 36, 37]) may affect the geometry of the phase space differently, both inputs keep the RNN to be a autonomous dynamical system. Specifically, varying the additive parameters within the input vector can modify the phase space geometry by moving the fixed point locations and changing the size of their domains of attraction. Additionally, in cases where the parameter values are close to *bifurcation values* (see [40]) which can lead to changes in the fixed points’ stability or to their disappearance.

If the dependence on the parameters of the right-hand side of the equations of motion is smooth, their SSMs will also display smooth dependency on parameters [28], [29]. However, whenever a fixed point disappears due to a bifurcation, we can no longer rely on the existence of its SSMs as invariant manifolds for the system.

In our examples, fixed points generally do not undergo bifurcations that lead to their disappearance, except in the context of the decision-making RNN when the relevant sensory inputs are varied. In that case, though, the reduced dynamics are carried by a Normally Hyperbolic Invariant Manifold (NHIM) [41] [42]. Therefore, the NHIMs theory and the asymptotic properties, uniqueness, and smoothness of unstable manifolds guarantee the persistence of a 1D invariant slow manifold for the whole parameter range considered, even if the fixed points undergo a cascade of saddle-node bifurcations (for details, see the Supplementary Methods S4).

In all the other cases, we can rely on the smooth dependence of SSMs on parameters to find parameter-dependent reduced-order models for the RNN dynamics. In fact, we can extend the phase space of parameter-dependent dynamical systems 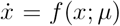, with *µ* ∈ ℝ^*p*^ the parameter vector, by treating the parameters as *dummy variables* for the system, which have trivial dynamics 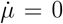. We can therefore find polynomial parameter-dependent expansions for SSMs and reduced-order models by solving the invariance equations in the extended phase space, including the parameter directions in the spectral subspaces.

On the other hand, to examine the impact of bounded noise on the dynamics of recurrent neural networks (RNNs), we apply an extension of SSM theory to nonautonomous systems subjected to weak aperiodic forcing, as discussed in [49]. We treat uniformly bounded noise as a form of discontinuous-in-time, weak forcing and derive time-dependent SSMs and reduced-order models that capture the noisy dynamics (refer to Supplementary Methods S6 for the general discussion on nonautonomous SSMs). These models will maintain the same phase space geometry as the autonomous system at any given moment, but they will evolve over time due to the influence of noisy perturbations.

In particular, fixed points for the autonomous system perturb, in the noisy setting, into noisy *anchor trajectories* with the same stability properties. We can think of such a trajectory as the perturbation of the line that would be present in the extended phase space with coordinates *X* = (*x, t*) ∈ ℝ^*N* +1^ with dynamics 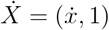, *L* = {(*x, t*) ∈ ℝ^*N* +1^ |*x* = 0, *t* ≥ 0}.

Any autonomous SSM perturbs into a time-dependent SSM that evolves over time along the anchor trajectory perturbing from the fixed point the autonomous SSM was attached to. The time-dependent SSM can be seen as the result of perturbing with additive, small noise terms the autonomous SSM in the extended phase space ℝ^*N* +1^ with coordinates (*y, t*). A more detailed discussion and the calculations specific for the noisy RNN systems can be found in the supplementary methods S6.1 and S6.2.

#### 4.2 Data-Driven Computations of SSMs

As we have seen, SSMs can be found locally as invariant graphs over their tangent spaces at the fixed point and approximated through Taylor expansions. This makes it possible to use a simple data-driven algorithm to obtain low-dimensional, predictive models for high-dimensional, nonlinear systems for which the equations of motions are not known (see [35]).The data-driven algorithm that finds SSMs and reduced models on them, SSMLearn, is open-source and available at https://github.com/haller-group/SSMLearn.

In this study, we combine the equation and data-driven approaches, since we use simulated RNN trajectories to find optimal polynomial SSMs and reducedorder model approximations through SSMLearn.

More specifically, SSMLearn is an algorithm designed to find the optimal polynomial coefficient matrix *H* for the SSM equations

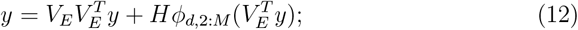

where *d* is the dimension of the SSM and *M* is the order of approximation of the polynomial expansion. The function *ϕ*_*d*,2:*M*_ (*q*) outputs the vector of all *m*_*M*_ monomials from order two up to order *M* given a *d*-vector *q, H* is the *N* × *m*_*M*_ coefficient matrix, 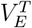 is the projection matrix to the spectral subspace,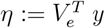 is the *d*-dimensional coordinate vector projected to the spectral subspace.

The optimization problem for *H*, and possibly the *N* × *d*-matrix *V*_*e*_, is formulated as minimizing a quadratic cost function that measures the distance between train trajectories and the SSM. When an estimate of the spectral subspace is already available (*V*_*e*_ is known), the optimization problem is convex.

It is then possible to compute the polynomial expansion for the reduced equations of motions 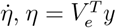. In particular, SSMLearn finds the optimal polynomial coefficient matrix *W*_*r*_ for the equation

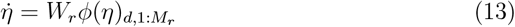

minimizing the distance between trajectories *η*(*t*) and the train trajectories projected to the spectral subspace. Finally, full-order trajectories can be obtained by lifting the reduced-order trajectories through the SSM equations.

### 5 The Vanilla RNN Models

The RNNs described in [39] and [18] are trained to obtain specific readout dynamics associated with behavioral experiments. These RNNs are designed to produce the correct output values based on a given set of input parameters, enabling them to replicate the target behaviors exhibited by trained animals. In this context, the optimal RNN parameters are identified to accurately reproduce a *N*_*o*_-dimensional readout function **z** that depends on the specific task.

The network units are coupled through a connectivity matrix **W** and have nonlinear dynamics due to a bounded, smooth, nonlinear activation function. They have equations

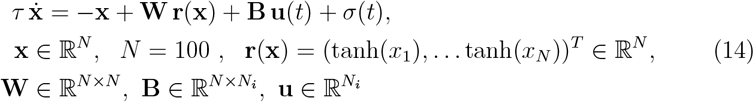

where **u** is the input vector of dimension *N*_*i*_ and *σ* ≈ *A N*(0, 1) ∈ ℝ ^*N*^ is a vector with entries drawn at each time step from a Normal distribution with zero mean and unit standard deviation^4^.

The matrices **W, B** and 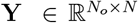 (*N*_*o*_ is the readout vector dimension) are selected by training to produce a *read-out function z*(*t*) = ⟨**Y**, **r**(**x**(*t*)) ⟩ with specific dynamics adapted to each task.

We first remove the noisy components from equations (14) and choose constant input vectors (which, in the experiments in [13], [39], [18], are only varied among trials). We then vary the input parameters to obtain manifolds and reduced ODEs that explicitly depend on them and describe the parametric changes to the geometry of the phase space. Finally, we insert noise back into the equation to obtain a noisy, time-dependent model.

#### 5.1 RNN for Context-Dependent Decision-Making

We now describe the specific features of the *Vanilla RNN* described in Eq. 14 that adapt it to the context-dependent decision-making task of Mante et al. in [13].

In the experiment reported in [13], monkeys were trained to distinguish between the direction of motion (left or right) or the color (red or green) of a random-dot display based on the given context cue. Meanwhile, extracellular responses from the monkey’s neurons were recorded in and around the frontal eye field and analyzed in the space of the first three principal components.

At each trial, neurons in the RNN receive two independent sensory inputs, each with a positive or negative value, that mimic evidence for motion and color in a single random dot stimulus, and the context inputs (see Fig. 5, right) indicate which of the two sensory inputs will determine the sign of the final scalar readout *z* ∈ ℝ (*N*_*o*_ = 1).

**Fig. 5:**
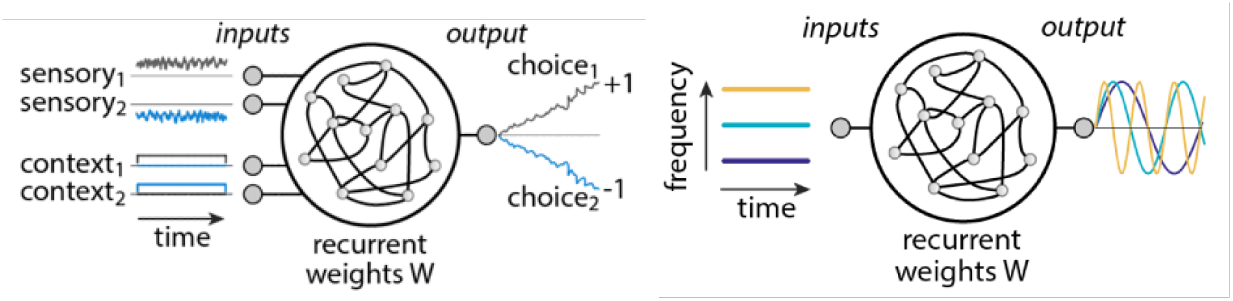
Pictorial representation of the context-dependent task performed by the RNN in [13] and [39] (right). Pictorial representation of the sine-wave generator RNN in [39] (left).

The 100D network evolves according to Eq. (14), with a 4D (*N*_*i*_ = 4) input vector **u** = (*u*_1_ *u*_2_ *u*_3_ *u*_4_)^*T*^ that accounts for both context and sensory stimuli in the experiment. In particular, its first two entries *u*_1_ = *s*_1_ and *u*_2_ = *s*_2_ (represent sensory stimuli (color and direction of motion, respectively). The network was trained on the set of values {− 0.15, − 0.036, − 0.009, 0.009, 0.036, 0.15}. The last two entries in the input vector *u*_3_ and *u*_4_ are one-hot-encoded and represent context 1 when *u*_3_ = *c*_1_ = 1 and *u*_4_ = *c*_2_ = 0, and context 2 when *u*_4_ = *c*_2_ = 1 and *u*_3_ = *c*_1_ = 0.

The input vector **u** is piecewise constant during one trial: in the first part of the trial, before the *burnlength time T*_*B*_, the sensory inputs are switched off *s*_1_ = *s*_2_ = 0 and only one of the context inputs is on. During this period, the target readout *z*^*\**^(*t*) = *const*. Instead, when *t > T*_*B*_, the sensory inputs are switched on *u*_1_ = *s*_1_, *u*_2_ = *s*_2_ and the sensory integration process begins. The target readout is a scalar function converging to either +1 or −1, depending on the selected sensory input.

We will treat this discontinuous-in-time dynamical system as the union of two separate systems, one for each time interval.

#### 5.2 Sine-Wave Generator RNN

The sine-wave generator network is a 100D Vanilla RNN with equations of motions of the same structure as in Eq. (14). The behavior implemented in this case is not linked to a specific experiment, but is rather a general model for oscillatory responses, which are of interest in the study of muscle contraction (see [39]).

The input *u* is, in this case, a scalar (*N*_*i*_ = 1), whose values represent the desired oscillation frequencies. The network is trained so that the scalar readout *z*(*t*) = ⟨**Y, r**(**x**(*t*) ⟩ evolves in time as a sine-wave with the frequency indicated by the scalar input (see Fig. 5, left).

#### 5.3 Multitasking RNN: Memory-Pro Task

The multitasking RNN implemented in [18] is

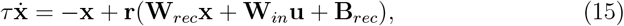

a variant of the Vanilla RNNs described in Eq. 14 with **x** ∈ ℝ^256^. The matrices **W**_*rec*_, **W**_*in*_, **B**_*rec*_ are trained so that the RNN can perform multiple tasks, depending on the values of the 20D input vector **u**. The one-hot-encoded last fifteen entries *u*_*i*_, *i* = 6, …, 20 indicate which task the network has to perform in each trial. The first entry of the input vector plays the role of a fixation input, taking values 0 or 1 depending on whether the network, similarly to animals in an experiment, has to respond to sensory inputs or not. Finally, the two 2D angular sensory inputs are represented in the entries *u*_2_ = cos *θ*_1_, *u*_3_ = sin *θ*_1_, *u*_4_ = cos *θ*_2_, and *u*_5_ = sin *θ*_2_.

When the fixation input is off (*u*_1_ = 0), the network has to respond according to the angle indicated by the stimulus that is relevant to the current task. The threedimensional readout vector **z** (*N*_*o*_ = 3) encodes whether the network is fixating or not in its first entry *z*_1_ = 1 or *z*_1_ = 0, and for the output angle through the values of *z*_2_ = cos *θ*_*out*_ and *z*_3_ = sin *θ*_*out*_.

In a Memory-Pro task, the RNN is exposed to piecewise constant changes in the input vector. In the context period [*t*_0_, *t*_*s*_], the sensory inputs are off, while the fixation and context inputs are on. Then, in the stimulus period *t*_*s*_, *t*_*m*_, sensory inputs are switched on, while all the other inputs remain constant. Sensory inputs are switched off again in the memory period [*t*_*m*_, *t*_*r*_], after which the fixation stimulus is switched off, and the network has to output the correct angle *θ*_*out*_ (see Fig. 6).

**Fig. 6:**
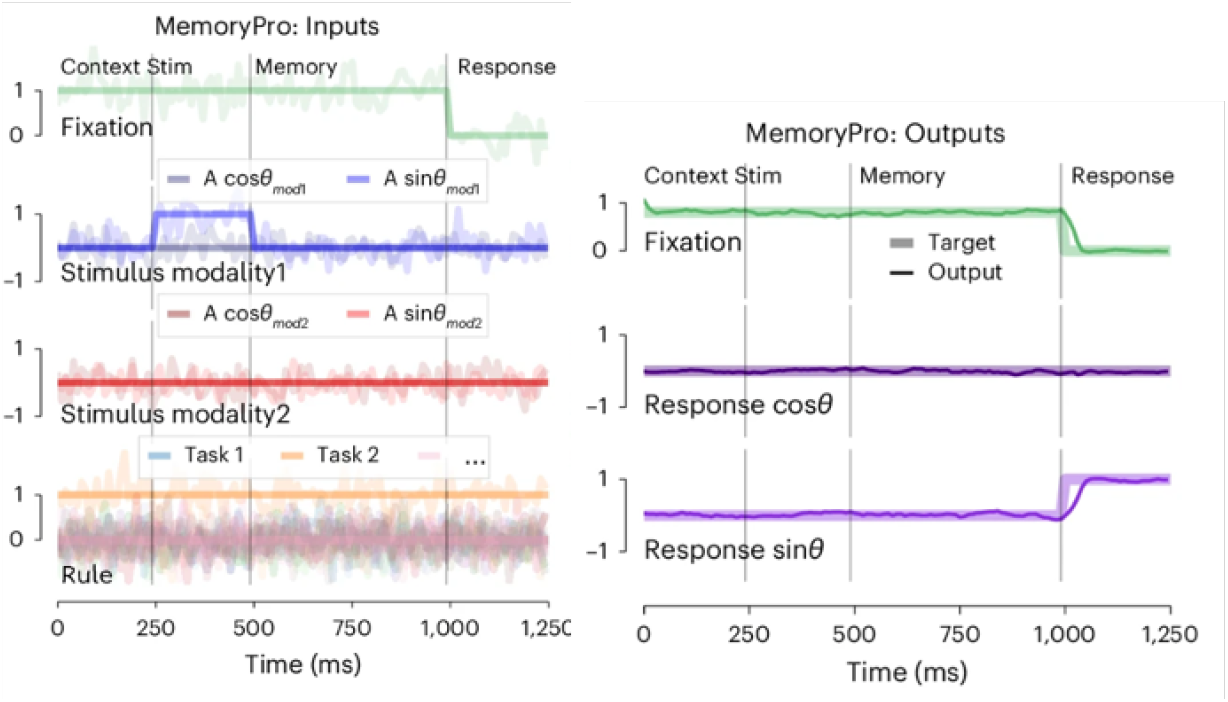
Pictorial representation of the inputs (right) and outputs (left) to the multitasking RNN in [18] performing a Memory-Pro task.

This task is designed to understand possible dynamical motifs that underlie working memory in animals. In our analysis, we will focus on the configuration during context and memory periods, with context and fixation input on and sensory inputs off.

## Footnotes

4 In our treatment, we will reject drawings outside ± 3 standard deviations from the mean to make the Gaussian noise bounded.

