## Supplementary Material for "Data-Driven Reduced Modeling of Recurrent Neural Networks"

### Supplementary Information

#### S1 Context-Dependent Decision-Making RNN: Details on Reduced Dynamics

We begin our analysis of the dynamical system in Eq. (14) by removing the noise components and fixing the sensory and context inputs. In particular, for the computations shown in this section, we choose $u_1 = s_1 = u_2 = s_2 = 0$ and $u_3 = c_1 = 0$, $u_4 = c_2 = 1$ when $t < T_B$ (sensory inputs are off and we choose context 2) and we switch on the sensory inputs to $u_1 = s_1 = 0.036$, $u_2 = s_2 = 0.15$ when $t > T_B$.

We obtain two autonomous dynamical systems with constant inputs, one for $t < T_B$ and the other for $t > T_B$. When $t < T_B$, we expect to find evidence of the set of *slow points* in the phase space found by Mante et al. in [13]. When $T > T_B$, we expect this line of slow points to disappear and to uncover some mechanism that can account for the binary choice performed by the network according to the input vector's entries. In both cases, a reduced-order model on a one-dimensional SSM can explain the dynamics of the network.

For the sensory-inputs-off period $t < T_B$, we find a stable fixed point in the phase space region explored by trajectories starting from initial conditions within a ball $B_{5\delta}$, $\delta = 0.1$. We are interested in this specific region of the phase space, since the network in [39] and [13] was always initialized at zero, and the bounded noise contributions have maximal amplitude $3\delta$. The network was trained to obtain a constant readout dynamics, so convergence to a stable fixed point is a simple realization of this behavior. We also find the two additional stable fixed points corresponding to the two choices, which turn out to be present for all parameter values explored in the task (see Fig. S2), but trajectories do not explore their domains of attraction in the pre-stimulus period.

The linearized dynamics around the stable fixed point has eigenvalues with a large gap between the real part of the first eigenvalue $\lambda_1$ and all the real parts of the other eigenvalues, which are smaller. Based on this observation regarding the linearized dynamics, we can expect that, at the considered timescales, the dynamics will quickly decay to the (existing, unique and smooth) 1D slowest Spectral Submanifold tangent to $E_1 = \mathrm{span}\{\mathbf{e}_1\}$ (eigenvector of eigenvalue $\lambda_1$). We find the SSM pictured in Fig. S1 (left), and Fig. 1, at fifth order and the reduced-order model for the network dynamics on it through SSMLearn. In Fig. 1 (center left), we plot the right-hand side of the reduced 1D equations of motions up to the fifth order.

Trajectories quickly converge to the slowest SSM. Once they have approached the manifold, they slowly approach the stable fixed point. This creates the impression of a set of "slow points" moving along an invariant curve in the phase space, converging slowly to a stable fixed point. Therefore, the line attractor reported by Mante et al. in [13], is just the slowest SSM attached to the stable fixed point: it attracts nearby trajectories and drives slow dynamics due to the (infinite-time) convergence to the stable fixed point.

We now turn to the analysis of the sensory integration period $T > T_B$ and set $\mathbf{u} = (0.15, 0.15, 0, 1)^T$. We find one unstable fixed point and two stable fixed points, i.e., two-point attractors in the phase space. Trajectories with random initial conditions are more likely attracted to the fixed point with the larger domain of convergence.

The unstable fixed point has a single unstable eigenvalue, hence a 1D unstable manifold. Since the gap between the real parts of the unstable eigenvalue and stable eigenvalues is wide, we expect the unstable manifold to capture the core dynamics of the system.

We find the unstable manifold with SSMLearn up to order five (see Fig. 1, upper left, and Fig. S1, right), and the 1D reduced-order model on the unstable manifold (see Fig. 1, center left), through which it is possible to simulate test trajectories and predict the whole system's dynamics.

To complete our analysis, we add noise to the equations of motion (14). For illustrative purposes, we focus on the sensory input-off time interval when there is a stable fixed point in the explored region of the phase space, and the core dynamics are captured by a reduced-order model on the slowest SSM. We also choose context 2 in the input vector $\mathbf{u}$ to be consistent with the results above for the autonomous network.

We apply the results of [49] for weak, time-dependent, uniformly bounded, and possibly discontinuous-in-time forcing terms. With this aim, we treat the noise term $\sigma(t) \approx 0.1\, \mathcal{N}(0,1) \in \mathbb{R}^N$ as small, discontinuous bounded forcing that depends on time and is independent of the position in the phase space.

The stable fixed point $\mathbf{x}_0$ for the autonomous system perturbs into an attracting, uniformly bounded anchor trajectory. We approximate this anchor trajectory through the formal expansions derived in [49]. We stop at order two, since the resulting approximated trajectory at order one is already a good approximation of the attracting anchor trajectory that we could estimate by evolving an initial condition in $\mathbf{0}$ with the noisy equations of motions 14. The formulas for these calculations can be found in the Supplementary Materials S6.1.

Once we have a good approximation of the anchor trajectory, we can compute the aperiodic, time-dependent slowest Spectral Submanifold $\mathcal{W}(E_1, t)$ attached to $\mathbf{y}^*(t) = \mathbf{x}^*(t) - \mathbf{x}_0$ when the sensory inputs are off and tangent, for every fixed $t$ to the spectral subspace $E_1 = \mathrm{span}\{\mathbf{e}_1\}$ in $\mathbf{y}^*(t)$. We truncate the approximation for $\mathcal{W}(E_1, t)$ at order $M = 2$. The calculations for the first and second-order terms of the expansion can be found in the Supplementary Materials S6.2.

The perturbed dynamics consist of fast convergence to the (moving) slowest stable SSM followed by slow convergence to the moving anchor trajectory.

#### S2 Context-Dependent Decision-Making RNN: Parameter Dependent SSMs

We recall that the 4D input vector $u$ in the decision-making RNN of [13] has entries $u_1 = s_1$, $u_2 = s_2$ corresponding to sensory inputs, and one-hot-encoded entries $u_3 = c_1$, $u_4 = c_2$ playing the role of context inputs. Here, we study changes in the phase space geometry due to parametrically varying both context and sensory inputs, and highlight how the role of the input vector in changing the RNN dynamics can be explained, in this case, in terms of parameter-dependent variations in the phase space.

We will consider a 1D parameter $\mu$, corresponding to either context or relevant sensory/irrelevant sensory inputs, and compute the $\mu$-dependent 1D slow manifolds and reduced-order models by looking for the 2D smooth invariant manifold in the extended phase space $\mathbb{R}^{N+1}$ with coordinates $(\mathbf{y}, \mu)$, tangent to the two-dimensional plane spanned by $\mathbf{e}_1$ (the eigenvector corresponding to eigenvalue $\lambda_1$ as above) and the vector spanning the parameter direction $\mathbf{c}_{N+1}$ in the extended phase space, at the point $(\mathbf{0}, 0)$.

We seek a such inveriant manifolds as invariant graphs over the plane $P = \text{span}\{\mathbf{e}_1, \mathbf{c}_{N+1}\}$ with coordinates $\eta_1$ and $\mu$ through SSMLearn. We will then find the coefficients $b_{j,l}$ for the corresponding two-dimensional models on the SSMs, obtaining reduced-order, predictive models at some order of approximation $q$ in $\eta_1$ and $\mu$

$$\begin{cases} \dot{\eta_1} &= \sum_{j+l\leq q} b_{j,l}\; \eta_1^j\; \mu^{\;l} \;+\; \mathcal{O}(q) \\ \dot{\mu} &= 0 \end{cases} \tag{S1}$$

We wish to understand how the dynamics are modified by varying the input vector $\mathbf{u}$ to achieve the flexibility needed for a context-dependent decision-making task. We expect that varying context inputs $u_3 = c_1$ and $u_4 = c_2$ would lead to significant changes in the domains of attraction of the two fixed points, whenever the sensory inputs have different signs.

We fix the sensory inputs to $s_1 = -0.15$, $s_2 = 0.15$ and parametrically vary the context inputs through the scalar $k$, $c_1 = 1 - k$, $c_2 = k$, $k \in [0, 1]$. The number and stability of the fixed points remain unchanged for different values of $k$, but we are interested in how the the global phase space geometry changes with $k$, which depends on the locations of the boundaries of the domains of attraction of the two stable fixed points.

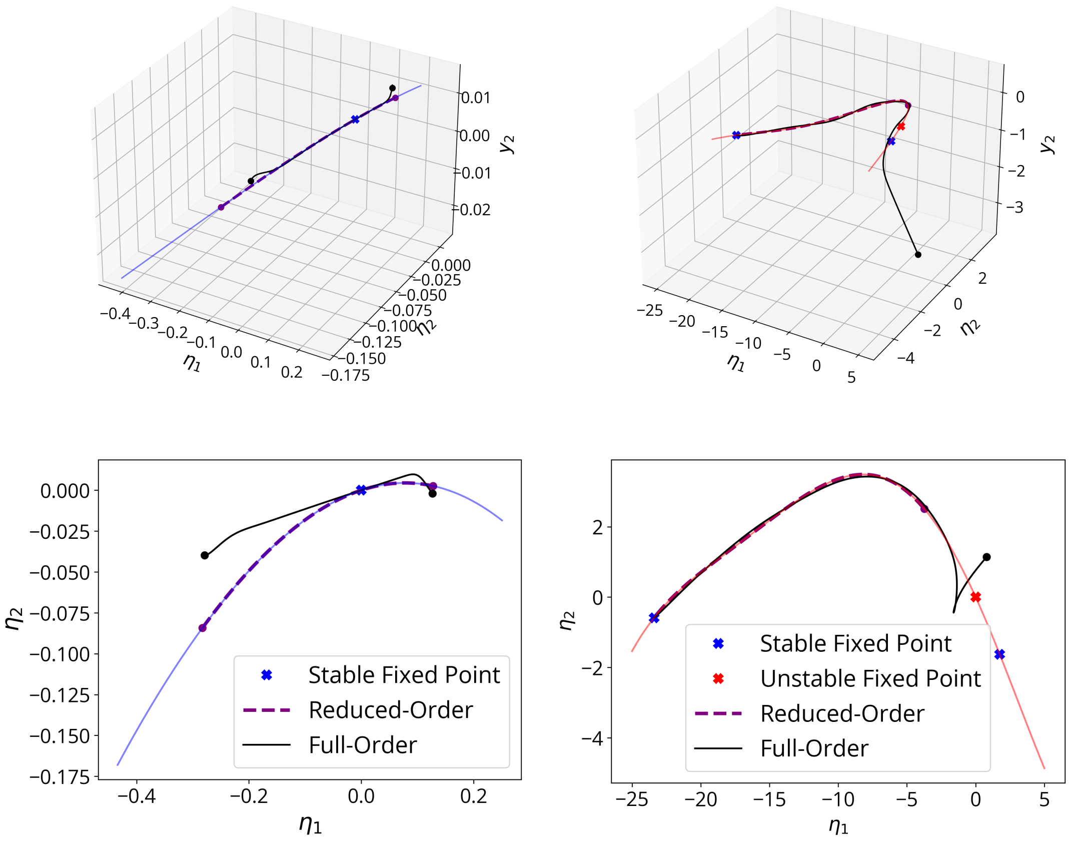


**Fig. S1:** (Top) The slowest SSM at order five (left, sensory inputs off, $s_1 = s_2 = 0$, context 2 $c_2 = 1,\ c_1 = 0$) and the unstable manifold of the unstable fixed point at order five (right, sensory inputs on $s_1 = 0.036, s_2 = 0.15$) and test full-order and reduced-order trajectories plotted with coordinates $(\eta_1, \eta_2, y_2)$, where $\eta_1, \eta_2$ parametrize the spectral subspace $E_2$ (corresponding to the first two Principal Components) and the $y_i$, $i = 1, \ldots, 100$, are coordinates for the 100D phase space $\mathbb{R}^N$, where the origin is at the anchor point. (Bottom) Manifolds projected on the spectral subspace $E_2$ of the unstable fixed point, spanned by the eigenvectors corresponding to its slowest eigenvalues (one stable and one unstable), with coordinates $(\eta_1, \eta_2)$, corresponding to the first two Principal Components space.

The 2D invariant manifold in the extended phase space $\mathbb{R}^{N+1}$ with coordinates $(\mathbf{y}, \bar{k})$, $\bar{k} = k - k_0$, $k_0 = 0.5$, exists and is smooth in $\mathbf{y}$ and $k$, for every value of $k \in [0, 1]$, due to the smooth dependence of the SSMs on the parameters, when the fixed points they are attached to are robust. Hence, we look for this invariant manifold as a graph over the extended spectral subspace $P$ through SSMLearn. We plot the slices of the extended phase space for different values of $k$ of the approximation of this manifold at order five in Fig. S3 (upper left), and the whole invariant manifold in the extended phase space in Fig. S3 (upper right). We find the corresponding reduced model at order nine as in Eq. S1, with $\mu = \bar{k}$ and $q = 9$.

We plot the parameter-dependent right-hand side in Fig. S3 (upper center). We deduce from the 1D model the widths of the domains of attraction of the two stable fixed points in the reduced model by looking at the location of the unstable fixed point that acts as a 0-dimensional boundary of such domains. These widths vary as expected for the network trained for the context-dependent task, since initial conditions around zero switch the fixed point they convergence to for $k = 0.5$.

Next, we study the effects of varying sensory inputs on the system's dynamics. First, we fix the context inputs at $c_1 = 0$, $c_2 = 1$ (context 2) and the select sensory input $s_2 = 0.036$, while we vary the sensory input $s_1$ in the range $[-0.036, 0.036]$. Since the network must ignore the sensory clues coming from the input $s_1$, we expect that the network dynamics will look the same when $s_1$ varies in the considered range.

Since the fixed point configuration is unchanged in the considered range of $s_1$, we can look for the 2D invariant manifold (which exists and is smooth by the smooth dependence of unstable manifolds on parameters) in the extended phase space $\mathbb{R}^{N+1}$ with coordinates $(\mathbf{y}, \bar{s}_1)$, $\bar{s}_1 = s_1 - s_{1,0}$, $s_{1,0} = 0$. The slices of this manifold for fixed $s_1$ correspond to the unstable manifolds of the unstable fixed points. We find an approximation of order five of the global manifold that yields a manifold fitting error of order $10^{-4}$.

In Fig. S3 (bottom left), we plot the resulting 1D, parameter-dependent manifolds obtained for fixed values of the parameter $s_1$ in the above equation, while on the bottom left we plot the global invariant manifold in the extended phase space. We also obtain the reduced-order model at order nine in $\eta_1$ and $s_1$ as in Eq. S1 with $\mu = s_1$ and $q = 9$ whose right-hand sides for the different values of $s_1$ are plotted in Fig.S3 (bottom center).

As expected, the phase space geometry does not undergo significant changes when varying the sensory input $s_1$, if context two is selected. The parameter-dependent reduced-order model on the $s_1$-dependent unstable manifolds correctly captures this phenomenon.

Finally, we study the consequences of parametric changes in sensory input 2 when context 2 is selected, and the first sensory input $s_1$ is kept fixed ($s_1 = 0.036$)

in the range for $s_2$ selected by Mante et al. in [13], $s_2 \in [-0.15, 0.15]$.

We find the system's fixed points for varying values of $s_2$ within the specified range using a numerical solver and observe that, even though the two stable fixed points corresponding to the two choices remain fixed, the number and location of the other fixed points change with the parameter. In particular, four saddle-node bifurcations occur within a small interval around $s_2 = 0$, as illustrated in the bifurcation diagram in Fig. S2 (right).

It is through this cascade of saddle-node bifurcations that the system adapts to favor the convergence to the stable fixed point that yields the correct readout (positive for positive values of $s_2$ and vice versa) along the branches of the 1D unstable manifold of an unstable fixed point.

If we take negative values of $s_2$, the unstable manifold that carries the reduced dynamics of the RNN is unique and normally hyperbolic. We show in Appendix S4 that even when the associated unstable fixed point (the bottom-most in Fig. S2, right) disappears, a 1D invariant slow manifold persists throughout the entire parameter range and continues to carry the reduced dynamics of the network.

However, since the unstable fixed point to which we attach the SSM does not persist along the whole $s_2$ range, it is not possible to find a single parameter-dependent reduced model for the dynamics through SSMLearn. Nevertheless, for each value of $s_2$, we can find a 1D reduced model for the observed dynamics on the 1D slow SSM, and we can describe the network dynamics globally for the whole parameter range in the extended phase space (see Fig. S2, left).

In particular, we already knew (see Fig. 1, bottom) that when $s_2$ is positive, the 1D unstable manifold of an unstable fixed point carries the decision-making reduced dynamics, leading convergence to the stable fixed point that corresponds to a positive readout. On the other hand, when $s_2$ is negative, the unstable manifold of the bottom-most unstable fixed point in Fig. S2 (right) arising from a saddle-node bifurcation when $s_2 \approx 0$ leads convergence to the fixed point corresponding to a negative value of the readout. These unstable manifolds correspond to line attractors when sensory inputs are on in [13].

When $s_2 \approx 0$, instead, the stable fixed point appeared in the critical $s_2$ interval around zero (where the four saddle-node bifurcations take place) attracts all the trajectories in a ball $B_{5\delta}$, $\delta = 0.1$ around zero, carrying a 1D slowest SSM (line attractor when sensory inputs are off in [13]).

These results yield a comprehensive interpretation of the system's behavior as governed by a global, 1D slow manifold structure that smoothly deforms for input parameter changes and whose internal dynamical structure adapts to consistently lead to convergence to the correct fixed point indicated by the inputs.

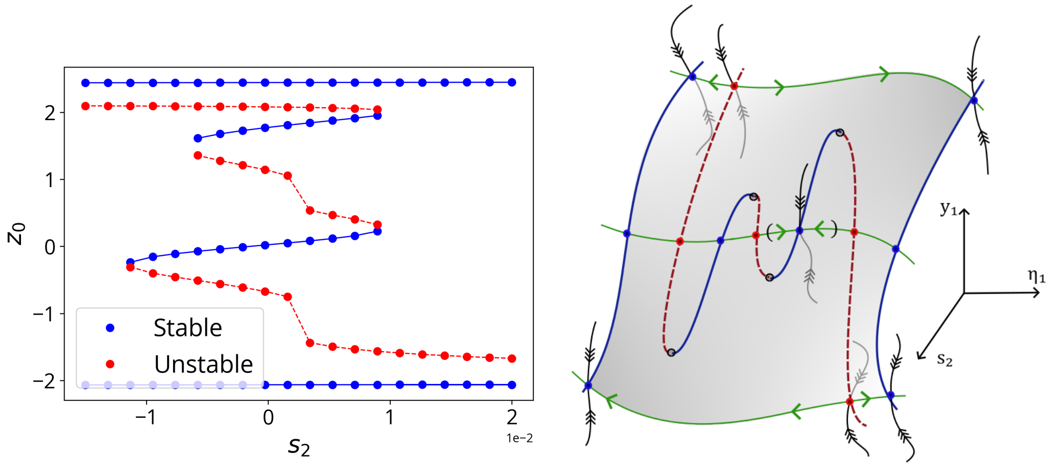


**Fig. S2:** (Left) Bifurcation diagram for the fixed points of the RNN, when sensory input 2 $s_2$ (selected by context input 2) is varied in the range $s_2 \in [-0.15, 0.15]$ and sensory input 1 is kept fixed at $s_1 = 0.036$. Four saddle-node bifurcations occur in the plotted critical parameter range $[-0.015, 0.020]$ around zero: two create a pair of fixed points, one stable and one unstable, and two annihilate one of such pairs. Through this cascade of saddle-node bifurcations, the system adapts by converging to a fixed point that yields the correct readout: negative when $s_2 < 0$, and positive when $s_2 > 0$. We plot a one-dimensional readout defined by $z_0 = \langle \mathbf{Y}, \mathbf{x}_0 \rangle$. (Right) Sketch of the robust, normally hyperbolic 2D slow manifold in the extended phase space, the global invariant manifold that carries the reduced dynamics of the RNN. When $s_2 < 0$ ($s_2 > 0$), the negative-readout (positive-readout) fixed point has the largest domain of attraction, and trajectories converge to it along a branch of the unstable manifold of the unstable fixed point that acts as a boundary. When $s_2 = 0$, the stable fixed point around zero appeared via a saddle-node bifurcation attracts trajectories along its 1D slowest SSM (in round brackets).

#### S3 Finite-Time Lyapunov Exponent (FTLE)

To estimate the sensitivity of RNN trajectories to initial conditions, we compute the *Finite-Time Lyapunov Exponent (FTLE)*. Given the flow map $F_{t_0}^t(x_0)$ for the system $\dot{x} = f(x, t)$, the FTLE quantifies the maximal rate of trajectory separation:

$$\mathrm{FTLE}_{t_0}^t(x_0) = \frac{1}{2|t - t_0|} \log \lambda_{\max}(C_{t_0}^t(x_0)), \tag{S2}$$

where $C_{t_0}^t = [DF_{t_0}^t(x_0)]^T DF_{t_0}^t(x_0)$ is the Cauchy-Green strain tensor (see [54] for more details).

We evaluate the FTLE over a grid of initial conditions on the two-dimensional spectral subspace $E_2 = \mathrm{span}\{\mathbf{e}_1, \mathbf{e}_2\}$, aligned via a linear change of basis around the unstable fixed point. The transformed system is:

$$\dot{\tilde{y}} = -\tilde{\mathbf{y}} - T\mathbf{x_0} + T\mathbf{W}\tanh(T^{-1}\tilde{\mathbf{y}} + \mathbf{x_0}) + T\mathbf{B}\ \mathbf{u} \tag{S3}$$

where $\tilde{y} \in \mathbb{R}^N$ are the new coordinates and $T$ the transformation. Simulating the dynamics allows us to compute $D_{\tilde{y}_1, \tilde{y}_2} F^t(\tilde{\mathbf{y}}_0)$ for any point $\tilde{\mathbf{y}}_0$ on the plane numerically by finite differences and, from it, the restricted FTLE.

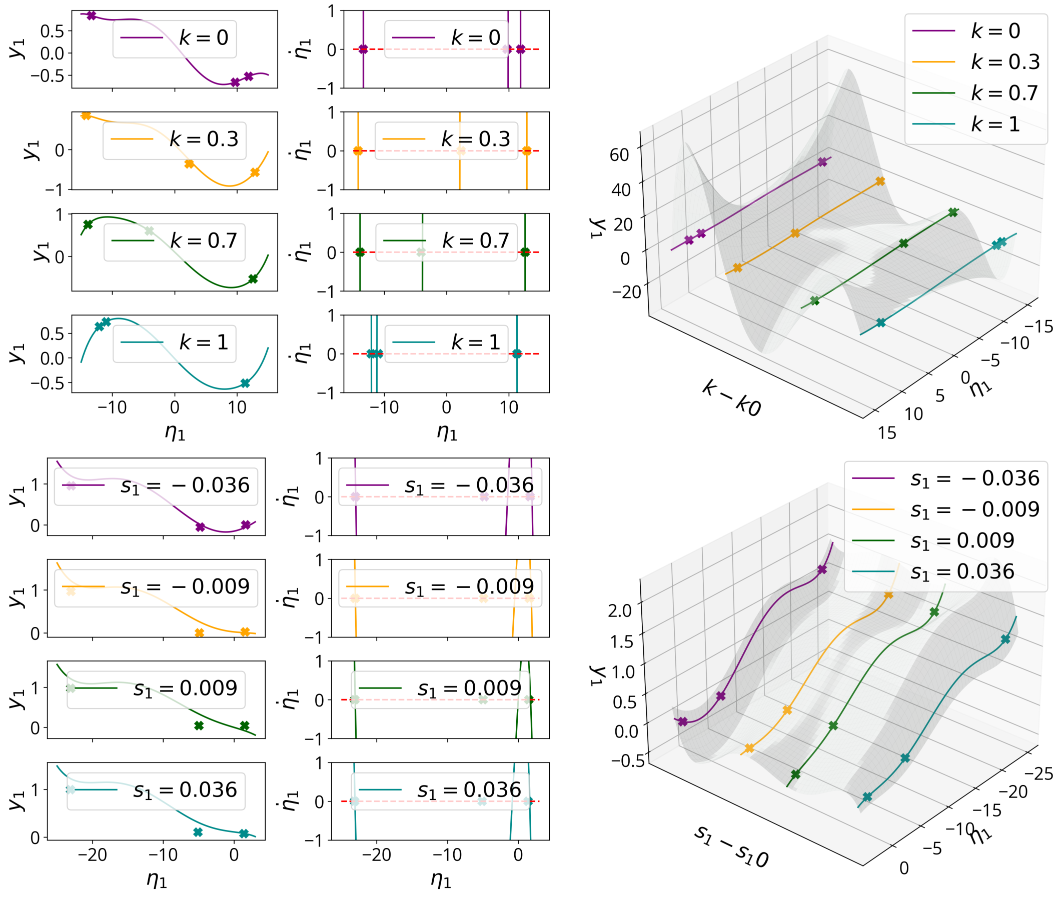


**Fig. S3:** (Left) Parameter-dependent SSMs yielding the unstable manifolds for different values of $k$ (top), with $c_1 = k$ and $c_2 = 1 - k$, when $s_1 = -0.15$ and $s_2 = 0.15$, and for different values of $s_1$ (bottom), where $s_2 = 0.15$ and context 2 is selected. The manifolds are plotted in coordinates $(\eta_1, y_1, )$. The symbols $x$ indicate fixed points of the full model on the manifolds (two stable and one unstable in the middle, acting as boundary of the domains of attractions of the other two). (Center) Graphs of the right-hand sides of the parameter-dependent reduced-order models on the unstable manifolds for different values of $k$ (top) parametrizing the context inputs, $c_1 = k$ and $c_2 = 1 - k$, and $s_1$ (sensory input 1). The symbols x indicate fixed points of the full-order model, to be compared to points where $\dot{\eta}_1 = 0$. We note how the domains of attraction of the two stable fixed points change parametrically with the context input $k$ as expected. When varying $s_1$, instead, the reduced dynamics do not vary, as expected (bottom). (Right) Visualization of the 2D slow manifolds in the extended phase space in coordinates $(\eta_1, \mu, y_1)$, where $\mu = \bar{k} = k - k_0$ (top) and $\mu = \bar{s}_1 = s_1 - s_{1,0}$ (bottom).

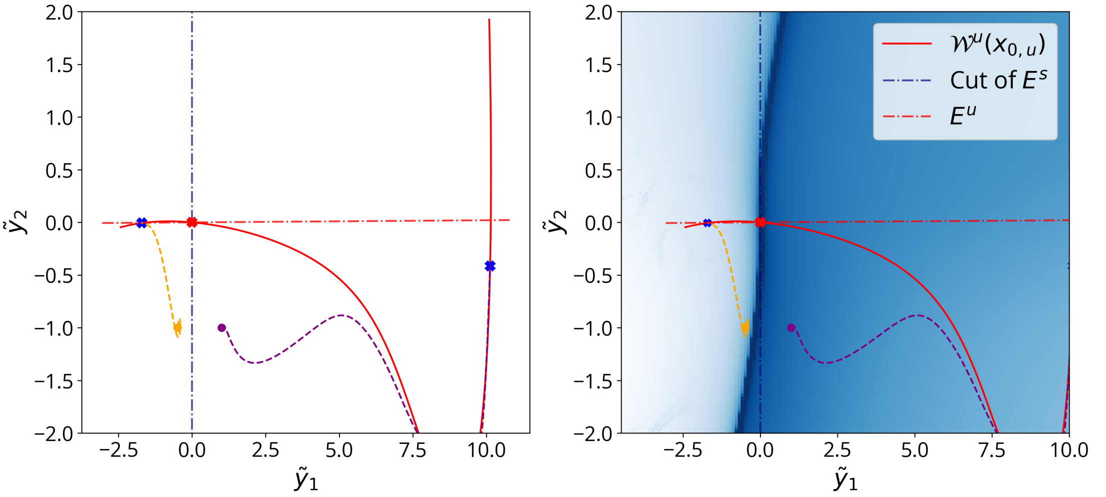


**Fig. S4:** (Left) Projection of the dynamics around the unstable fixed point on the spectral subspace $E_2$. We plot the projected intersections of the unstable and stable subspaces ($E^S$ and $E^U$, respectively). (Right) FTLE calculated on the chosen two-dimensional plane $P = E_2$. Darker values in the picture on the left indicate higher values of the FTLE. Regions of high values of the FTLE indicate the estimated boundaries of the domains of attraction of the two fixed points. As expected, the cut of the stable manifold $\mathcal{W}^S$ on the plane $P$ is tangent to the stable subspace $E^S$. Trajectories with initial conditions belonging to the two different regions converge to different fixed points.

FTLE ridges, local maxima of the FTLE field (see [55]), indicate the boundaries between the domains of attraction of the two stable fixed points (see Fig. S4). These ridges correspond to intersections of the stable manifold $W^S(x_0)$ of the unstable fixed point with the selected subspace.

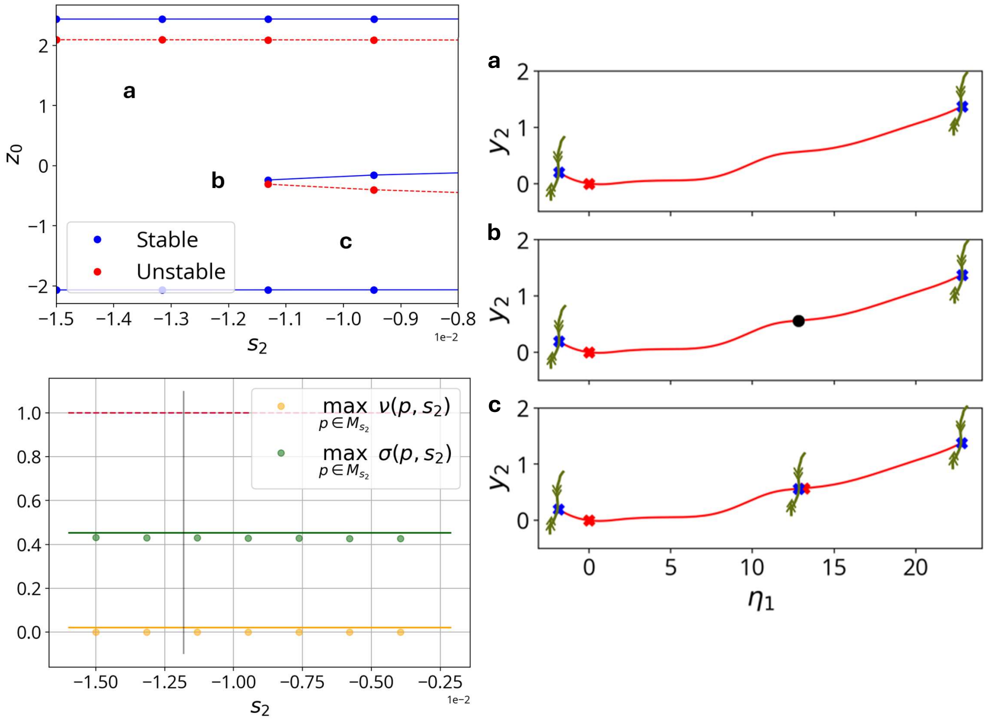


**Fig. S5:** (Upper left and right) Bifurcation diagram in the restricted parameter range where the first saddle-node bifurcation happens along a 1D center manifold, and the corresponding normally attracting, unique invariant manifolds in cases (a), (b) and (c). This is the bifurcation responsible for the appearance of the unstable fixed point from which we construct the unstable manifold for positive values of $s_2$ (see Fig. S2 and Fig. 1, bottom). (Bottom right) Supremum of type numbers for $\alpha$-limit sets of the invariant manifold over the $s_2$ range. Dots: numerical values; lines: upper bounds. Bounds below 1 indicate normal hyperbolicity throughout.

### Supplementary Methods

#### S4 Persistence of the Slow Manifold for the Context-Dependent Decision-Making task

In the context-dependent RNN of [13], the reduced dynamics lie on 1D unstable manifolds (when sensory inputs are on) or on the slowest SSM of a stable fixed point (when sensory inputs are off, $s_1 = s_2 = 0$) (see 5.1, 1). These manifolds coincide with the line attractor slow dynamics identified in [13].

To study the persistence of such a 1D slow manifold for the entire range of input parameters explored in the task, we consider $\epsilon$-parameter changes with $0 < \epsilon \ll 1$. Our goal is to determine whether the invariant manifold $\mathcal{W}_0$ of the system at $\epsilon = 0$ persists as a $C^\rho$-smooth manifold $\mathcal{W}_\epsilon$ that remains $\mathcal{O}(\epsilon)$-close to the original one.

Smooth dependence of SSMs on parameters generally suffices to state this persistence result, as long as the fixed points to which they are attached remain hyperbolic during parameter changes. In this case, though, when we vary the relevant sensory input for the task from negative to positive values ($s_1$ for context 1 and $s_2$ for context 2), we observe four saddle-node bifurcations shown in Fig. S2 (left). These bifurcations cause the unstable fixed points that we utilize to describe the reduced dynamics for both positive and negative sensory inputs through their unstable manifolds (see Fig. S2 and Fig. 1, bottom) to lose their hyperbolicity and disappear.

Nevertheless, a few simple observations let us establish a persistence result for this problem. We start by looking at negative relevant sensory input values ($s_2$ for context $c_2$) to focus on the parameter range where the first of such bifurcations happens, as shown in the bifurcation diagram in Fig. S5.

In region (a), there are two stable and one unstable fixed points. To show there is a unique, attracting invariant manifold in the RNN phase space, we can use the asymptotic property of the unstable manifold of the latter, which is defined as the set of points that converge to the unstable fixed point asymptotically in backward time. Therefore, we can extend the local unstable manifold by forward-time integration of initial conditions along the unstable eigenspace in the proximity of the fixed point.

Fig. S5 (right, a) shows the results of forward integration, through which we can conclude that there exists a 1D attracting invariant manifold $\mathcal{W}_0$ composed of the two stable fixed points, the unstable fixed point, and the two branches of its unstable manifold. This manifold is unique in class $C^0$ and, since the convergence to the stable fixed points along the unstable manifold must align with their slowest eigenvectors (the line spanned by $e_1$), $\mathcal{W}_0$ is also at least of class $C^1$. Moreover, it is compact with an invariant boundary, with the two stable fixed points serving as its closure.

We can now invoke Fenichel's results on the persistence of compact, overflowing-invariant manifolds (see [41], [42]). These results require the manifold to be normally hyperbolic: the rate of contraction or expansion in directions transverse to the manifold must dominate the dynamics along the manifold itself by some integer $\rho$. This smoothing property of the flow ensures persistence and determines the degree of smoothness of the perturbed manifold $\mathcal{W}_\epsilon$.

While an exposition of the theory of Normally Hyperbolic Invariant Manifolds (NHIMs) is beyond our scope, we briefly state the main result: a compact, $C^r$, $\rho$-NHIM will persist under small $C^1$ $\mathcal{O}(\epsilon)$-perturbations as a nearby invariant manifold of differentiability class $C^{\max\{\mathrm{r},\rho\}}$ for the perturbed flow (see [41, 42] for rigorous statements and proofs).

We now wish to show that the invariant manifold $\mathcal{W}_0$ is an attracting NHIM. To do so, we need to verify the definition of normal hyperbolicity through the exponential rate of normal attraction and tangential contraction of the linearized dynamics along the manifold. The asymptotic properties of the linearized flow are quantified through the *Lyapunov-type numbers*, for which we again refer to [41] and [42]. Since these numbers are defined by looking at asymptotic properties of trajectories on the manifold in backward time, to determine the normal hyperbolicity of a manifold, it is sufficient to calculate them on its $\alpha$-limit sets, i.e., the sets to which trajectories on the manifold converge in forward time.

In this case, the only limit sets are the fixed points. In particular, we have that the invariant manifold $\mathcal{W}_0$ is $\rho$-normally attracting if, given the ordered eigenvalues of the linearized dynamics around a fixed point $p$, $\lambda_1 \geq \lambda_2 \geq \cdots \geq \lambda_N$

1. The first Lyapunov type number quantifying the asymptotic rate of normal attraction $\nu(p) = e^{\lambda_2} < 1$ for all the fixed points $p$ in $W_0$ ($\mathcal{W}_0$ attracts nearby trajectories)

2. The second Lyapunov type number $\sigma(p) = \frac{\lambda_1}{\lambda_2} < \frac{1}{r}$, for some integer $r > 1$, for all the fixed points on $\mathcal{W}_0$ (the normal decay on $\mathcal{W}_0$ dominates over the tangential compression along $\mathcal{W}_0$ (if present))

As shown in Fig. S5 (bottom left), not only are the requirements on the Lyapunov type numbers satisfied, but there also exists a uniform bound along the whole considered parameter range.

Therefore, by Fenichel's theorem, we can conclude that there exists a semi-open interval of parameters $\epsilon \in [0, \bar{\epsilon})$ (which parametrizes some semi-open interval for $s_2$ $[-0.15, s_2^*)$) along which this manifold exists, is at least $C^1$ and is normally attracting. We now wish to prove that we can extend this persistence result to include the parameter value $\bar{s}_2$ corresponding to the bifurcation point (b) in Fig. S5, which is the only value where we cannot use the uniqueness and asymptotic properties of the unstable manifold to state the existence, uniqueness and smoothness result for the invariant manifold.

The uniform boundedness of the Lyapunov type numbers below one is a necessary condition to conclude this result (see [56]). As stated in [57], though, it is not sufficient to determine the existence of a $C^1$-smooth manifold at the boundary of the parameter range. In fact, this manifold might lose normal hyperbolicity by losing $C^1$ smoothness.

The complete argument goes as follows. Let $A = \{\epsilon \in [0, \bar{\epsilon}] : \forall \epsilon' \in [0, \epsilon]$, there exists a NHIM$\mathcal{W}_{\epsilon'}\}$. If $A \neq \emptyset$, $A$ open and $A$ closed, then $\bar{\epsilon} \in A$, and the result follows. We know that $A$ is nonempty, since $\mathcal{W}(0)$ is a NHIM, and the fact that $A$ is open follows from Fenichel's theorem. To show that $A$ is closed, we need to show that if $\epsilon^*$ is the supremum of $A$, then $\epsilon^* \in A$. In other words, we need to find an argument for the existence of a $C^1$ invariant manifold for $\epsilon^*$.

In our setting, the only value in $A$ we need to check is the bifurcation value $\epsilon^* = \bar{\epsilon}$. Since we know from calculating the Jacobian that the saddle-node bifurcation that gives rise to the pair of unstable and stable fixed points has a one-dimensional center manifold, and the center manifold is $C^r$-smooth for $r \in \mathbb{N}$, the existence of a $C^1$ invariant manifold is guaranteed at the bifurcation value. The manifold is normally hyperbolic, as shown in Fig. S5.

To show the existence of an attracting slow manifold in the extended phase space, describing the reduced dynamics of the context-dependent decision-making RNN for the whole considered $s_2$ range as in Fig. S2 (right), it remains to show that these normally hyperbolic manifolds we proved to exist are also unique.

We already know that in region (a), the manifold is unique by the uniqueness of the unstable manifold. In region (c), we can similarly invoke the definition of unstable manifold and continue the 1D branches emanating from each unstable fixed point by forward integration of trajectories along their unstable direction. The resulting invariant manifold, composed of three stable fixed points, two unstable fixed points, and the four branches of their unstable manifolds, is therefore also unique. The uniqueness at the bifurcation point follows from the uniqueness of the limit.

#### S5 Normal Hyperbolicity of the Heteroclinic Structure for the Memory-Pro Task

We aim to establish the normal hyperbolicity of the heteroclinic manifold $\mathcal{W}_h$ in the 2D reduced-order model of the multitasking RNN presented in [18], which performs the Memory-Pro task.

The manifold $\mathcal{W}_h$ consists of a stable fixed point, an unstable fixed point, and the two branches of the unstable manifold connecting them. This set forms a compact, boundaryless, and invariant manifold by construction.

As shown in Figure 4, trajectories initially converge onto the two-dimensional unstable manifold that contains $\mathcal{W}_h$, and subsequently contract rapidly onto $\mathcal{W}_h$ itself, ultimately reaching the stable fixed point. To verify normal hyperbolicity, we evaluate the Fenichel-type numbers $\nu(p)$ and $\sigma(p)$ for all $p \in \mathcal{W}_h$, characterizing the asymptotic behavior as $t \to +\infty$.

The only limit sets within $\mathcal{W}_h$ are the two fixed points. The unstable one has a 1D unstable manifold and N-1 decaying directions in the linearized dynamics. The linearized dynamics around the stable fixed point have the first two eigenvalues satisfying $\mathrm{Re}\lambda_{2,s} < \mathrm{Re}\lambda_{1,s} < 0$. The eigenvector corresponding to $\lambda_2$ is tangent to $\mathcal{W}_h$, while the one associated with $\lambda_1$ is transverse. We can calculate:

- $\max_{p\in\mathcal{W}_h} \nu(p) = \max\left\{e^{\lambda_{2,s}},\ e^{\lambda_{2,u}}\right\} = 0.0156 < 1$ (normal attraction rate)

- $\max_{p\in\mathcal{W}_h} \sigma(p) = \frac{\lambda_{1,s}}{\lambda_{2,s}} = \frac{-0.0496}{-5.0213} < \frac{1}{101}$

Hence, $\mathcal{W}_h$ is $\rho$-normally attracting with $\rho = 101$.

### S6 Nonautonomous Spectral Submanifolds (SSMs)

SSM theory extends to nonautonomous systems with periodic, quasiperiodic, weak aperiodic, slow aperiodic and even random forcing. We briefly summarize the key results and refer the reader to [28, 49, 50] for formal statements, assumptions, and proofs.

For periodically or quasiperiodically forced systems of the form

$$\dot{x} = Ax + f_0(x) + \epsilon f_1(x, \Omega t; \epsilon), \quad x \in \mathbb{R}^N,\ \Omega \in \mathbb{R}^k, \tag{S4}$$

with smooth nonlinearities $f_0 = \mathcal{O}(\|x\|^2)$ and $f_1$ periodic in each $\Omega_i t$, fixed points of the unforced system perturb into periodic (or quasiperiodic) solutions known as *Nonlinear Normal Modes (NNMs)*. Invariant manifolds dependent on time, non-autonomous SSMs, exist in their vicinity under non-resonance conditions [28].

In the RNN analysis, the only time-dependent input is bounded aperiodic forcing, in the form of drawings from a bounded Gaussian distribution. For this setting, we consider systems of the form

$$\dot{x} = Ax + f_0(x) + f_1(x, t), \tag{S5}$$

where $f_1$ is uniformly bounded in time but not necessarily continuous. If the unforced system has a hyperbolic fixed point, this point perturbs into a uniformly bounded, compact trajectory called an *anchor trajectory* under specific uniform boundedness conditions on the time-dependent forcing $f_1$ [49]. SSMs can then be constructed along this trajectory, provided a spectral subspace $E$ is $\rho$-normally hyperbolic and satisfies nonresonance conditions

$$\sum_{j=1}^{N} m_j \mathrm{Re}\lambda_j \neq \mathrm{Re}\lambda_k \ \ \forall k \in \{1, \ldots, n\}, \quad m_j \in \mathbb{N},\ \sum_{i=1}^{N} m_j \geq 2. \tag{S6}$$

The key result is that under these assumptions, a unique time-dependent SSM $\mathcal{W}_E(x^*(t))$ exists, is tangent to $E$ at each time step, and can be computed as the uniformly bounded solution of a time-dependent nonlinear PDE, even for larger forcing amplitudes than predicted by theory. These manifolds enable model reduction and capture the slow nonlinear dynamics of recurrent neural networks (RNNs) under single realizations of bounded Gaussian noise in the form of weak aperiodic forcing.

Formal expansions for both anchor trajectories and time-dependent SSMs are derived in [49]. In our work, specific expansions for RNN dynamics are presented in S6.1 and S6.2.

#### S6.1 Anchor Trajectory Calculations

First, we want to compute anchor trajectories $\mathbf{x}^*(t)$ perturbing from fixed points when small-amplitude discontinuous forcing is added to the equations of the autonomous RNNs.

We have the rescaled equations of motions for the Vanilla RNNs

$$\mathbf{x}' = \frac{d\mathbf{x}}{dt'} = \frac{1}{\tau}\frac{d\mathbf{x}}{dt} = \frac{1}{\tau}\Big(-\mathbf{x} + \mathbf{W}\,\mathbf{r}(\mathbf{x}) + \mathbf{B}\,\mathbf{u}\Big) \tag{S7}$$

We change coordinates to $\mathbf{y} = \mathbf{x} - \mathbf{x_0}$ so that the fixed points we want to attach SSMs to are in zero. In the following, we change notation from $\mathbf{y}'$ to $\dot{\mathbf{y}}$ to indicate the derivative with respect to the rescaled time $\frac{t}{\tau}$, and we denote such rescaled time with $t$. Hence,

$$\begin{aligned} \dot{\mathbf{y}} &= \mathbf{A}\,(\mathbf{y} - \mathbf{x_0}) + \mathbf{f}(\mathbf{y}, t), \\ \mathbf{f}(\mathbf{y}, t) &= \mathbf{f}_0(\mathbf{y}) + \epsilon \mathbf{f}_1(\mathbf{y}, t) + \mathbf{B}\mathbf{u}, \quad \mathbf{f}_0(\mathbf{y}) = O(|\mathbf{y}|^2), \end{aligned} \tag{S8}$$

with

$$\begin{aligned} \mathbf{A} &= -\mathbf{I} + \begin{pmatrix} \frac{w_{11}}{\cosh^2(x_{1,0})} & \cdots & \cdots & \frac{w_{1N}}{\cosh^2(x_{N,0})} \\ \vdots & \vdots & \vdots & \vdots \\ \frac{w_{N1}}{\cosh^2(x_{1,0})} & \cdots & \cdots & \frac{w_{NN}}{\cosh^2(x_{N,0})} \end{pmatrix}, \\ \mathbf{f}_0(\mathbf{y}) &= \mathbf{W}\tanh(\mathbf{y} - \mathbf{x_0}) - \begin{pmatrix} \frac{w_{11}}{\cosh^2(x_{1,0})} & \cdots & \cdots & \frac{w_{1N}}{\cosh^2(x_{N,0})} \\ \vdots & \vdots & \vdots & \vdots \\ \frac{w_{N1}}{\cosh^2(x_{1,0})} & \cdots & \cdots & \frac{w_{NN}}{\cosh^2(x_{N,0})} \end{pmatrix} (\mathbf{y} - \mathbf{x}_0), \\ \mathbf{f}_1(\mathbf{y}, t) &= \mathbf{f}_1(t) = \sigma(t), \end{aligned}$$

and $\sigma(t)$ is drawn at each time step from a bounded standard normal distribution.

We use the formal expansions in [49] to approximate anchor trajectories perturbing from fixed points through $\mathbf{y}^*(t) = \sum_{\nu=1}^{M} \epsilon^\nu \mathbf{y}_\nu(t) + o(|\epsilon \mathbf{f}_1|_U^M)$, assuming it exists.

We calculate the expansion terms $\mathbf{y}_\nu(t)$ up to order $M = 3$, and perform (discrete) numerical integration to obtain these contributions numerically at each time step. For the first-order term, given $\mathbf{A}$ with $d_u$ unstable eigenvalues, we have

$$\mathbf{y}_1(t) = \int_{-\infty}^{t} \mathbf{T} e^{\begin{pmatrix} \mathbf{0} & \mathbf{0} \\ \mathbf{0} & \mathbf{A}_s \end{pmatrix}(t-\tau)} \mathbf{T}^{-1}\mathbf{f}_1(\tau)\, d\tau \; - \int_{t}^{+\infty} \mathbf{T} e^{\begin{pmatrix} \mathbf{A}_u & \mathbf{0} \\ \mathbf{0} & \mathbf{0} \end{pmatrix}(t-\tau)} \mathbf{T}^{-1}\mathbf{f}_1(\tau)\, d\tau,$$
$$\mathbf{A} = \mathbf{T}\begin{pmatrix} \mathbf{A}_u & \mathbf{0} \\ \mathbf{0} & \mathbf{A}_s \end{pmatrix}\mathbf{T}^{-1} \quad \mathbf{A}_u = \begin{pmatrix} \lambda_1 & \mathbf{0} & \dots \\ \mathbf{0} & \ddots & \vdots \\ \vdots & \dots & \lambda_{d_u} \end{pmatrix} \quad \mathbf{A}_u = \begin{pmatrix} \lambda_{d_u+1} & \mathbf{0} & \dots \\ \mathbf{0} & \ddots & \vdots \\ \vdots & \dots & \lambda_N \end{pmatrix}, \tag{S9}$$

We write the expressions for the second and third-order contributions

$$\mathbf{y}_2(t) = \int_{-\infty}^{t} \mathbf{T} e^{\begin{pmatrix} \mathbf{0} & \mathbf{0} \\ \mathbf{0} & \mathbf{A}_s \end{pmatrix}(t-\tau)} \mathbf{T}^{-1}[\,\frac{1}{2}\partial_{\mathbf{y}}^2\mathbf{f_0}(\mathbf{0}) \otimes \mathbf{y}_1(\tau) \otimes \mathbf{y}_1(\tau)]\, d\tau$$
$$- \int_{t}^{+\infty} \mathbf{T} e^{\begin{pmatrix} \mathbf{A}_u & \mathbf{0} \\ \mathbf{0} & \mathbf{0} \end{pmatrix}(t-\tau)} \mathbf{T}^{-1}[\,\frac{1}{2}\partial_{\mathbf{y}}^2\mathbf{f}_0(\mathbf{0}) \otimes \mathbf{y}_1(\tau) \otimes \mathbf{y}_1(\tau)]\, d\tau$$
$$\mathbf{y}_3(t) = \int_{-\infty}^{t} \mathbf{T} e^{\begin{pmatrix} \mathbf{0} & \mathbf{0} \\ \mathbf{0} & \mathbf{A}_s \end{pmatrix}(t-\tau)} \mathbf{T}^{-1}\Big[\frac{1}{6}\partial_{\mathbf{y}}^3\mathbf{f}_0(\mathbf{0}) \otimes \mathbf{y}_1(\tau) \otimes \mathbf{y}_1(\tau) \otimes \mathbf{y}_1(\tau)$$
$$+ \frac{1}{2}D_{\mathbf{y}}^2\mathbf{f}_0(\mathbf{0}) \otimes \mathbf{y}_1(\tau) \otimes \mathbf{y}_2(\tau)\Big]\, d\tau$$
$$- \int_{t}^{+\infty} \mathbf{T} e^{\begin{pmatrix} \mathbf{A}_u & \mathbf{0} \\ \mathbf{0} & \mathbf{0} \end{pmatrix}(t-\tau)} \mathbf{T}^{-1}\Big[\frac{1}{6}\partial_{\mathbf{y}}^3\mathbf{f}_0(\mathbf{0}) \otimes \mathbf{y}_1(\tau) \otimes \mathbf{y}_1(\tau) \otimes \mathbf{y}_1(\tau)$$
$$+ \frac{1}{2}D_{\mathbf{y}}^2\mathbf{f}_0(\mathbf{0}) \otimes \mathbf{y}_1(\tau) \otimes \mathbf{y}_2(\tau)\Big]\, d\tau,$$

where

$$\mathbf{f}_0(\mathbf{y}) = \begin{pmatrix} w_{11}\tanh(y_1 - x_{1,0}) + ... - (\frac{w_{11}}{\cosh^2(x_{1,0})}(y_1 - x_{1,0}) + ... + \frac{w_{1N}}{\cosh^2(x_{N,0})}(y_N - x_{N,0})) \\ \vdots \\ w_{N1}\tanh(y_1 - x_{1,0}) + ... - (\frac{w_{N1}}{\cosh^2(x_{1,0})}(y_1 - x_{1,0}) + ... + \frac{w_{NN}}{\cosh^2(x_{N,0}}(y_N - x_{N,0})) \end{pmatrix},$$
$$\partial_{\mathbf{y}}\mathbf{f}_0(\mathbf{x}_0) = \mathbf{0},$$
$$\partial_{\mathbf{y}}\mathbf{f}_0(\mathbf{y}) = \begin{pmatrix} \frac{w_{11}}{\cosh^2(y_1 - x_{1,0})} & \dots & \frac{w_{1N}}{\cosh^2(y_N - x_{N,0})} \\ \vdots & \vdots & \vdots \\ \frac{w_{11}}{\cosh^2(y_1 - x_{1,0})} & \dots & \frac{w_{1N}}{\cosh^2(y_N - x_{N,0})} \end{pmatrix} - \begin{pmatrix} \frac{w_{11}}{\cosh^2(x_1^0)} & \dots & \frac{w_{1N}}{\cosh^2(x_N^0)} \\ \vdots & \vdots & \vdots \\ \frac{w_{11}}{\cosh^2(x_1^0)} & \dots & \frac{w_{1N}}{\cosh^2(x_N^0)} \end{pmatrix}.$$

Note that $\partial^2_{\mathbf{y}} \mathbf{f}_0(\mathbf{y})$ is a 3-tensor wherein each vector entry is a Hessian matrix:

$$\partial^2_{\mathbf{y}} \mathbf{f}_0(\mathbf{y}) = \begin{pmatrix} \begin{bmatrix} -\frac{2w_{11}\sinh(y_1 - x_{1,0})}{\cosh^3(y_1 - x_{1,0})} & & \\ & \ddots & \\ & & -\frac{2w_{1N}\sinh(y_N - x_{N,0})}{\cosh^3(y_N - x_{N,0})} \end{bmatrix} \\ \vdots \\ \begin{bmatrix} -\frac{2w_{N1}\sinh(y_1 - x_{1,0})}{\cosh^3(y_1 - x_{1,0})} & & \\ & \ddots & \\ & & -\frac{2w_{NN}\sinh(y_N - x_{N,0})}{\cosh^3(y_N - x_{N,0})} \end{bmatrix} \end{pmatrix}$$

Finally, $\partial^3_{\mathbf{y}} \mathbf{f}_0(\mathbf{y})$ is a 4-tensor (each $N$-vector entry is a $N \times N$ matrix of row $N$-vectors):

$$\partial^3_{\mathbf{y}} \mathbf{f}_0(\mathbf{y}) = \begin{pmatrix} \begin{bmatrix} \begin{bmatrix} w_{11} f^3(y_1 - x_{1,0}) & 0 & \dots & 0 \end{bmatrix} & & \\ & \ddots & \\ & & \begin{bmatrix} 0 & \dots & 0 & w_{1N} f^3(y_N - x_{N,0}) \end{bmatrix} \end{bmatrix} \\ \vdots \\ \begin{bmatrix} \begin{bmatrix} w_{N1} f^3(y_1 - x_{1,0}) & 0 & \dots & 0 \end{bmatrix} & & \\ & \ddots & \\ & & \begin{bmatrix} 0 & \dots & 0 & w_{NN} f^3(y_N - x_{N,0}) \end{bmatrix} \end{bmatrix} \end{pmatrix}$$

where

$$f^3(y_i - x_{i,0}) = \frac{4\sinh^2(y_i - x_{i,0})}{\cosh^4(y_i - x_{i,0})} - \frac{2}{\cosh^4(y_i - x_{i,0})} = \frac{4\sinh^2(y_i - x_{i,0}) - 2}{\cosh^4(y_i - x_{i,0})}$$

#### S6.2 Nonautonomous SSM Calculations

We calculate the aperiodic, time-dependent SSMs, $\mathcal{W}(E, t)$, attached to the anchor trajectories obtained above $\mathbf{y}^*(t) = \mathbf{x}^*(t) - \mathbf{x}_0$, perturbing from the autonomous SSMs, $\mathcal{W}(E)$, tangent to a spectral subspace $E = E_1 \oplus \cdots \oplus E_k$.

Let $\mathbf{P} = [\mathbf{e}_1 \ \dots \ \mathbf{e}_N] \in \mathbb{C}^N$ be the matrix containing the complex eigenvectors corresponding to the ordered eigenvalues of $A$, and change coordinates to

$$\begin{pmatrix} \mathbf{u} \\ \mathbf{v} \end{pmatrix} = \mathbf{P}^{-1}(\mathbf{x} - \mathbf{x}^*_\epsilon) \quad \Longrightarrow \quad \begin{pmatrix} \dot{\mathbf{u}} \\ \dot{\mathbf{v}} \end{pmatrix} = \begin{pmatrix} \mathbf{\Lambda} & 0 \\ 0 & \mathbf{A}_{\mathbf{v}} \end{pmatrix} \begin{pmatrix} u \\ \mathbf{v} \end{pmatrix} + \hat{f}(u, \mathbf{v}, \epsilon; t)$$

with

$$\mathbf{u} \in \mathbb{R}^d,\ \mathbf{v} \in \mathbb{R}^{N-d}, \qquad \mathbf{\Lambda} = \begin{pmatrix} \lambda_1 & & \\ & \ddots & \\ & & \lambda_d \end{pmatrix}, \quad \mathbf{A_v} = \begin{pmatrix} \lambda_{d+1} & & \\ & \ddots & \\ & & \lambda_N \end{pmatrix},$$

$$\hat{f}(\mathbf{u}, \mathbf{v}, \epsilon; t) = \mathbf{P}^{-1}\Big[\mathbf{f_0}\Big(\mathbf{x}_\epsilon^* + \mathbf{P}\begin{pmatrix}\mathbf{u}\\ \mathbf{v}\end{pmatrix}\Big) + \mathbf{A}\mathbf{x}_\epsilon^* + \mathbf{A}\dot{\mathbf{x}}_\epsilon^* + \epsilon\, \mathbf{f_1}(t)\Big]$$

In the case of the decision-making RNN, the time-dependent SSM, $\mathcal{W}(E_1, t)$, perturbs from the autonomous slowest SSM $\mathcal{W}(E_1)$ attached to the stable fixed point in the pre-stimulus period.

The SSM, $\mathcal{W}(E_1, t)$, admits a formal asymptotic expansion

$$\mathcal{W}(E_1, t) = \Big\{(u, \mathbf{v}) \in U \subset \mathbb{R}\times\mathbb{R}^{N-1}\,;\, \mathbf{v} = \mathbf{h}_\epsilon(u, t) = \sum_{|(k,p)|\geq 1}^{M} \mathbf{h}^{kp}(t) u^k \epsilon^p + o(|u|^q, \epsilon^{M-q})\Big\}.$$

According to [49], we have

- $\mathbf{h}^{0,p} = 0$.
- $\mathbf{h}^{k,0}$ are the coefficients of the unperturbed manifold $\mathcal{W}(E_1)$ expansion over the space spanned by $\mathbf{e}_1$.
- The other coefficients $\mathbf{h}^{kp}$ can be obtained through the integral formula ([49])

$$\mathbf{h}^{kp} = \int_0^\infty \mathbf{G}_k(t-s)\, \mathbf{M}^{kp}(s, \mathbf{h}^{jm}(s))\, ds$$

  and $\mathbf{G}_k(t) = e^{(\mathbf{A_v} - k\,\lambda_1 \mathbf{I}_{N-1})t}$.

We stop at order $M = 2$, since first-order contributions already yield a good approximation of the time-dependent manifold. We have that

$$\begin{aligned}\mathcal{W}(E_1, t) = \Big\{(u, \mathbf{v}) \in U \subset \mathbb{R}^N\,;\, \mathbf{v} = \mathbf{h}_\epsilon(u, t) &= \sum_{|(k,p)|\geq 1}^{2} \mathbf{h}^{kp}(t) u^k \epsilon^p + o(|u|^q, \epsilon^{2-q}) \\ &= \mathbf{h}^{20} u^2 + \mathbf{h}^{11} u\ \epsilon + o(|u|^q, \epsilon^{2-q})\Big\},\end{aligned}$$

and

- The autonomous coefficients:
  - $\mathbf{h}^{10} = \mathbf{0}$.

– $\mathbf{h}^{20} = -\mathbf{A}_2^{-1} M^{20}(\mathbf{0}), \quad M^{20}(\mathbf{0}) = \left[ P^{-1} \frac{\partial^2 \mathbf{f}_0}{\partial u^2}(\mathbf{x}_0) P \begin{pmatrix} 1 \\ 0 \end{pmatrix} P \begin{pmatrix} 1 \\ 0 \end{pmatrix} \right] \Big|_{\mathbf{v}}.$

- $k = 1, p = 1$ means the anchor trajectory needs to be inserted at first order, and $j = 1$, $m = 0$:

$$M^{11}(t, \mathbf{0}) = \left[ P^{-1} \frac{\partial^2}{\partial^2 \mathbf{y}} \mathbf{f_0}(\mathbf{x_0}) \otimes \mathbf{y}_1(t) \otimes P \begin{pmatrix} 1 \\ 0 \end{pmatrix} \right] \Big|_{\mathbf{v}}.$$
